# Milo2.0 unlocks population genetic analyses of cell state abundance using a count-based mixed model

**DOI:** 10.1101/2023.11.08.566176

**Authors:** Kluzer Alice, John C Marioni, Michael D Morgan

## Abstract

Cell type proportions vary between individuals and are heritable, as demonstrated by statistical genetic analysis of flow cytometry data [1,2]. Higher-resolution cell states can be identified by single-cell RNA-sequencing, the scalability of which now makes it applicable to population-scale cohorts. However, the integration of statistical genetic analysis of cell states using cohort-scale single-cell data requires appropriate algorithms to account for and model the genetic relationships and complex batch-processing inherent to these studies. We describe Milo2.0, which enables the discovery of cell state quantitative trait loci (csQTL), scaling to millions of cells across hundreds of individuals. We identify > 500 csQTLs across peripheral blood immune states and investigate their relationship with the genetic regulation of gene expression. Moreover, we colocalise immune csQTLs with human traits and identify links between immune regulators, cell state abundance and immune-mediated disease.

## Introduction

Genetics seeks to link molecular and cellular regulatory programs to disease pathophysiology. Quantitative trait loci (QTL) mapping is often the method of choice to statistically link genetic variation to specific phenotypes, notably mRNA or protein abundance levels in defined tissues or peripheral blood sera. More recently, advances in single-cell omics methods have enabled the acquisition of single-cell data on population cohorts [3]. This has placed the large sample sizes required for quantitative trait locus (QTL) analyses within reach of single-cell studies (reviewed in [4]). Notably, the integration of population-scale single-cell gene expression profiling has led to single-cell expression QTL (eQTL) studies that first identify constituent cell types through iterative rounds of clustering, sub-clustering and merging before performing eQTL analyses within each cell type [5–9]. This workflow necessarily leads to the identification of genetic variation that alters cell-type composition, which has previously only been possible using cytometry-based approaches [1,2,10,11]. Therefore, single-cell population scale genetic analyses have the capacity to identify both eQTLs and cell-state QTLs (csQTLs) within the same study, which may help to better understand the genetic predisposition to complex diseases [1,10].

Genetic analysis of diverse populations can be biassed by the cryptic genetic relationships between individuals [12]. Mixed effect models and, more broadly, generalised linear mixed models (GLMMs), can model dependent observations, including cryptic genetic structure, to account for such relationships [12]. Indeed, GLMMs have become a popular tool in human genome wide association studies due to their flexibility and ability to account for genetic relationships and recent advances in computational scalability [13,14]. Despite their appeal for modelling of count data, which is common in genomics studies, negative binomial (NB)-GLMMs have not been widely adopted. One key reason for this is the need to marginalise over the random effects, which involves a high-dimensional integral that can be analytically intractable. Multiple approaches have been proposed to circumvent this problem, many of which approximate the marginal likelihood [15,16], or use slow numerical optimisation or Bayesian approaches to estimate the posterior distribution [17,18]. The latter involves using Markov Chain Monte Carlo, which is computationally burdensome, and infeasible for differential abundance (DA) testing of many thousands of traits; a variational inference approach has been described for linear mixed effect models, but not for NB-GLMMs [19].

Using single-cell cluster proportions or counts is an efficient approach for studying the genetic regulation of cell type composition when cell types are well delineated or easily separable [20,21]. However, the boundaries between different cell types or states are often blurred due to heterogeneity, or continuous biological processes such as development or differentiation. We recently developed a powerful computational method, Milo, to identify differentially abundant cell states in single-cell experiments without the need to define clusters and cell types [20]. Here, we combine this clustering-free approach with the power of new scalable algorithms and mixed models to unlock the genetic analysis of cell states in population-scale single cell studies.

## Results

### Improving Milo scalability using graph-based distance approximation

In the original Milo workflow a set of refined neighbourhoods on a nearest-neighbour graph are computed from a random subsample of single cells from the data set [20]. Milo then computes an archetypal (median) cell from the nearest neighbours of each sampled cell in the same reduced dimensional space used to construct the NN-graph and defines the nearest real cell in this space as the index cell of the neighbourhood. Consequently, highly connected cells are selected as the neighbourhood indices, which reduces the total number of index cells and hence neighbourhoods. However, the process requires computation of cell-to-cell distances within each neighbourhood, with a time complexity of 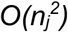 for the jth neighbourhood, which leads to poor scalability of the Milo workflow beyond ∼200,000 cells (Supplementary Figure 1A). We reasoned that the refined neighbourhood index cells all share a property - that of a high number of shared neighbours. Graph nodes that are highly connected will have a large number of 3-step cyclical connections, called triangles, and therefore have a large number of shared 1st order neighbours (Figure 1A). Computing the number of triangles in an undirected graph is equivalent to taking a 3-step random walk from each node, and is efficiently computable as the matrix trace of A^3^, where A is the adjacency matrix [22]. Importantly, selecting the cell with the largest number of triangles in each initial neighbourhood gives equivalent results to our original distance-based algorithm, recovering the same profile of nearest-neighbour relations using both methods (Figure 1B; Supplementary Figure 1B). Critically, the computational time required to compute triangles is 2 orders of magnitude faster than computing within neighbourhood distances and requires fewer computational resources (Figure 1C, Supplementary Figure 1C-E). We compared the performance of both refinement methods using 50,000-200,000 simulated single cells, as previously described [20]. While the complete Milo workflow required approximately 6 hours to complete using neighbourhood distances for 200,000 cells, the same analysis was completed in < 5 minutes using our new graph-based approximation.

**Figure 1.**
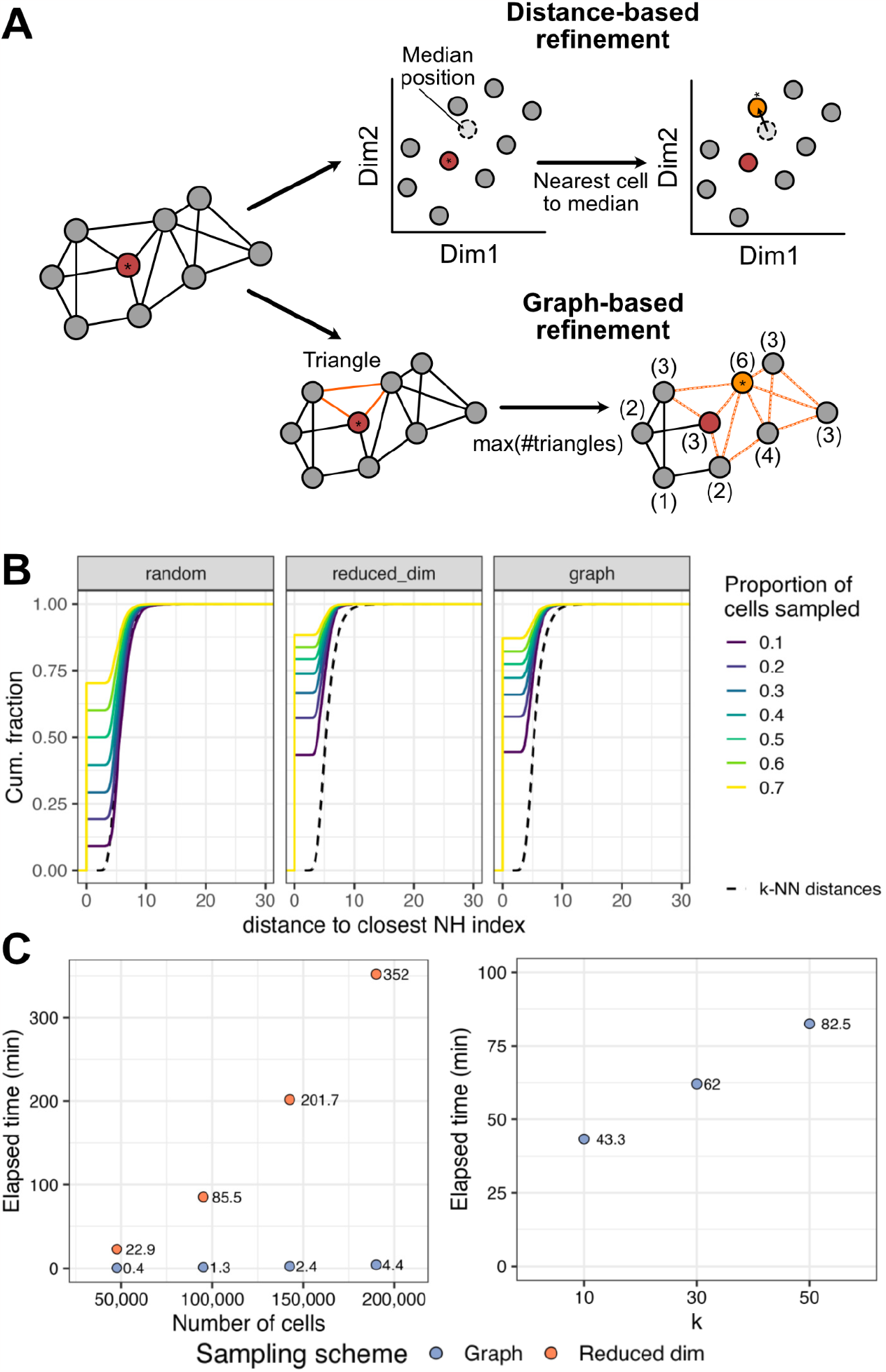
A graph-based neighbourhood refinement allows Milo2.0 to scale to population cohort-scale differential abundance analysis. (A) A schematic denoting the original (top) and graph-based (bottom) neighbourhood refinement algorithms. The asterisk denotes the chosen index cell. (B) Graph-based neighbourhood refinement achieves equivalent results to distance-based refinement. Shown are cumulative fractions of Euclidean distances to neighbourhood indices across random samplings with different initial proportions (colours). The black dotted line shows the distribution of kNN (k=30). (C) Run time of Milo analyses (y-axis) using distance-based (orange) or graph-based refinement (purple) on up to 200,000 simulated cells (left, k=30) and 1 million simulated cells (right) varying graph connectivity (x-axis). All times are shown in minutes.

From an experimental perspective, developments in scalability and multiplexing of single-cell experiment technology have enabled the collection of millions of cells across diverse donors, conditions and cell types [5,6,8]. The number of single-cells in a dataset is an important factor for Milo’s computational speed, as is the connectivity of the cells in the NN-graph, which is tunable using the parameter ‘k’ for the initial nearest neighbour searching. To test the impact of the number of graph connections, we applied Milo2.0 to a simulated data set of 1 million single cells and varied the value of ‘k’ used for the initial graph building step. As expected, the computational time increased with higher values of ‘k’ (Figure 1C), with the time taken for 1 million cells being approximately equivalent to 100,000 cells using our original graph-based refinement scheme.

Single cells can be members of multiple graph neighbourhoods. To account for this overlap we employ the spatial false discovery rate (FDR) to adjust for multiple hypothesis testing [20]. In our original implementation we used a weighted Benjamini-Hochberg procedure, using the inverse of the average cell-cell distance as the weighting for each neighbourhood. To fully dispense with cell-cell distance calculations, we reasoned that the number of cells shared between the neighbourhood in question and all other neighbourhoods would approximate the previously used weighting. This alternative measure has the advantage that it is trivially computable using the graph. Importantly, it achieves equivalent results to the distance-based correction, with no observed bias in the spatial FDR values, and with control of the FDR being maintained on simulated data (Supplementary Figure 1F).

In summary, the use of graph-based approximations to refine neighbourhoods and perform FDR correction achieves orders of magnitude improvement in computational efficiency while retaining the robustness of Milo differential abundance testing.

### A negative binomial mixed effect model extends the scope of Milo analyses

Milo performs DA hypothesis testing using the parameter estimates of log fold change from a negative binomial generalised linear model. This provides flexibility in terms of the different experimental designs that can be accommodated for DA testing with Milo. The growth of single-cell multi-tissue atlases and population-scale datasets motivates the need for even greater flexibility in experimental designs, particularly for those that include multiple observations from the same donor or organism. This is because GLMs assume that outcomes, such as gene expression or neighbourhood counts, are independent and identically distributed. However, multiple observations drawn from the same individual violate this assumption, which has been noted for single-cells from the same experimental sample [23]. A similar issue arises in genetic analyses where familial relationships and cryptic population structure can induce strong dependencies between individual observations [12]. To address this challenge, we extended Milo to incorporate random effect variables in a negative binomial generalised linear mixed model (NB-GLMM).

Motivated by the need for speed, we implement a pseudo-likelihood approximation of the model likelihood [24]. This is justified as (1) it allows the use of closed-form solutions for fixed and random effect weight estimates, (2) it is amenable to fast (quasi) Newton algorithms with near-quadratic convergence times, and (3) it allows for unbiased variance estimation using restricted maximum likelihood (REML). We have implemented Fisher scoring and Haseman-Elston regression (Methods) to allow for fast variance parameter estimation across a range of scenarios. Notably, this allows us the flexibility to use a pre-defined covariance matrix for the random effects, thus facilitating the incorporation of genetic effects into the analysis of cell state variation. While GLMMs can model both random intercepts and random slopes, the latter are less frequently used in biomedical research and require a more complex framework to solve; we therefore defer the implementation of random slopes models to future work. We implement the NB-GLMM framework in Milo2.0, which is written in the R programming language [25] that already includes several commonly-applied GLMMs that use different approximation approaches. To maintain cross-compatibility with the original Milo, we implemented computationally heavy aspects of the NB-GLMM in RCpp [26]. This provides interoperability and leverages the computational advantages of a statistically-typed language.

### Milo2.0 increases power to identify regulators of cell state variation

We previously demonstrated how adjusting for batch effects in Milo increased power and reduced false discoveries [20]. As single-cell population studies grow in size, accounting for batch effects with 10’s or more batches could have a significant negative impact on statistical power when modelled as a fixed effect as each batch uses an additional degree of freedom. To test this notion we performed DA testing using a single-cell data set of mock and influenza A virus (IAV) infected peripheral immune cells derived from a population cohort of European and African-similar ancestry individuals [9]. These data were generated in 30 batches and are thus an ideal test case for comparing the power gained by modelling batches as a random vs. fixed effect. We selected cells from uninfected samples, computed the kNN-graph from the batch-integrated dimensions (see Methods), and executed the complete Milo2.0 workflow to identify DA neighbourhoods between individuals from European or African ancestry-similar groups (Figure 2A-B). We computed population DA (popDA) neighbourhoods using our GLM with the batch variable as a fixed effect, and our NB-GLMM with batch as a random effect variable, which resulted in highly concordant log fold change estimates (Figure 2C), demonstrating the accuracy of our NB-GLMM in a real-world setting. Moreover, using a 10% FDR our NB-GLMM identified 312 popDA neighbourhoods between ancestry-similar groups distributed across a range of immune cell types (Figure 2D-F). By contrast, only 82 popDA neighbourhoods were identified with the GLM (Supplementary Figure 4), illustrating the large statistical power gains from modelling batch as a random effect. These gains were not specific to these data - we identified a similar pattern when using only the IAV infected samples (Supplementary Figure 5). We note that these findings cannot be explained by an elevated FDR in our NB-GLMM results, as random shuffling of ancestry labels and repeating the analysis 100 times did not lead to increased numbers of false discoveries (Supplementary Figure 5E-F). We likewise demonstrated the power gain using Milo2.0 compared to the GLM in an independent data set of T cells derived from COVID19 patients [27] (Supplementary Figure 6).

**Figure 2.**
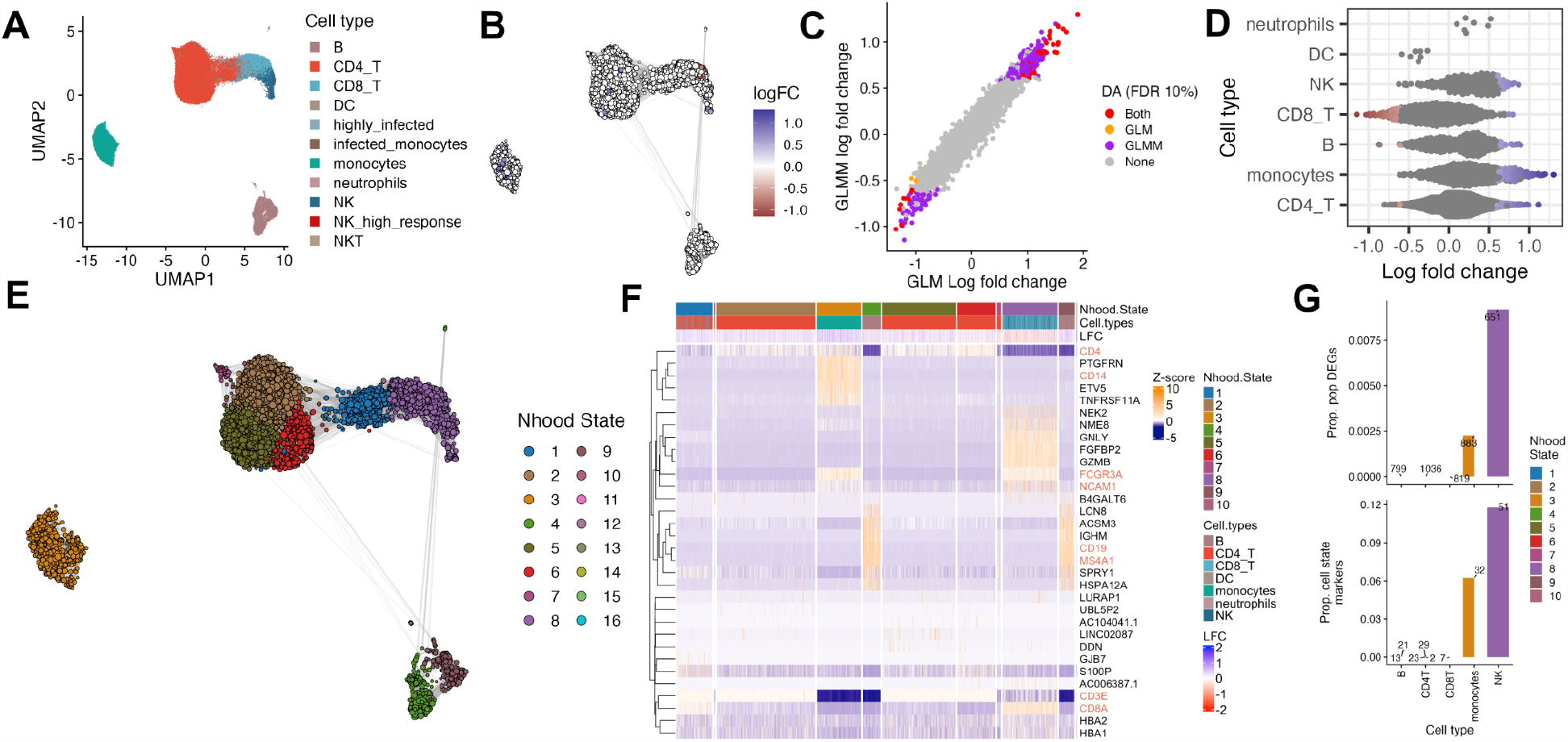
Milo2.0 increases power to identify regulators of cell state variation. (A) UMAP of *in vitro* mock infected peripheral immune single cells from Ref [9]. Points representing single-cells are coloured by their annotation in Randolph *et al*. (B) Milo neighbourhood graph illustrating differential abundant neighbourhoods overlaid on the UMAP embedding as in (A). Points are coloured by the log fold change between ancestry-similar groups. (C) GLM (x-axis) and GLMM (y-axis) log fold change estimates coloured by statistical significance threshold in the respective model (FDR ≤10%). (D) Beeswarm plot illustrating the relationship between DA neighbourhoods and annotated immune cell types from Randolph *et al*. Points are coloured by log fold change as in (C) y-axis. (E) Neighbourhood UMAP showing neighbourhood states (colours), where lines represent the number of shared cells between neighbourhoods. (F) Heatmap of popDA cell state markers overlaid with neighbourhood state and cell type information. Canonical cell type marker genes are highlighted in red (T cells: *CD4, CD3E, CD8A*, B cells: *CD19, MS4A1*, NK cells: *FCGR3A, NCAM1*, Monocytes: *CD14*). (G) Proportions of popDE genes (top) and popDA cell state markers (bottom) explained by the respective gene set.

At the cell type level, Randolph *et al*. previously identified 1949 population differentially expressed genes (popDE) between African and European ancestry-similar donors [9]. We investigated whether our popDA neighbourhoods could be explained by these cell type popDE genes. We first identified the gene expression markers for the popDA cell states in the mock and IAV infected samples separately (Figure 2E-F, Supplementary Figure 7A), then compared these with the cell type popDE gene set from [9]. Our analysis of the mock infected cells revealed that < 1% of cell type popDE genes were explained by popDA cell state markers (Figure 2G). In contrast, amongst the IAV infected cells >80% of monocyte popDE genes could be explained by markers for popDA cell states (Supplementary Figure 7B). However, the converse was not true as < 10% of popDA markers could be explained by monocyte popDE genes. Indeed, across all cell types the majority of popDA cell state marker genes were distinct from the cell type popDE genes indicating that Milo analysis provides complementary information on the ancestry differences underpinning immune system variation.

GLMMs model the variation between dependent observations. For example, they can be applied to experiments where multiple samples are derived from the same individual. Using the complete data from Randolph *et al*. [9] we performed DA testing between the mock and influenza infected samples while using the donor ID as a random effect in our GLMM. We compared our results to those using the GLM with no additional adjustment. We identified 1159 DA neighbourhoods from our GLMM and 1270 DA neighbourhoods with the GLM (FDR 10%). However, there was almost no concordance between the log fold changes between the two models (Spearman’s ρ=-0.02, Supplementary Figure 8A). We considered if this could be explained by an elevated FDR in the GLM caused by the violation of independence. Indeed, when we randomly shuffled the flu infection labels we found an elevated FDR in the GLM but not the GLMM (Supplementary Figure 8B-C). Together, these analyses demonstrate the extended capability of Milo2.0 to detect perturbed cell states and increased power gains for population cohort scale single-cell experiments.

### Milo2.0 unlocks population genetic analysis of cell state variation

Mixed effect models provide a powerful tool for adjusting for genetic relationships in trait-association studies [28]. Consequently, we reasoned that Milo2.0 would enable the identification of genetic variation that explains cell state variation across genetically diverse donors while accounting for this dependence. To demonstrate the feasibility of such an approach, we performed genome-wide association testing between common (MAF ≥ 15%) single nucleotide polymorphisms (SNPs) and neighbourhood abundance using the mock-infected samples from Randolph *et al*. [9]. At genome-wide significance (spatial FDR ≤ 10^-8^), our analysis identified 576 significant pairs of lead SNPs and neighbourhoods, which we term cell state quantitative trait loci (csQTLs, Figure 3A). These csQTLs were distributed across 335 distinct genetic loci (Supplementary Table 1) affecting monocytes and CD8+ T cells (Figure 3A). Importantly, csQTLs were not confounded with ancestry since csQTL and popDA neighbourhoods were distinct (Figure 3B).

**Figure 3.**
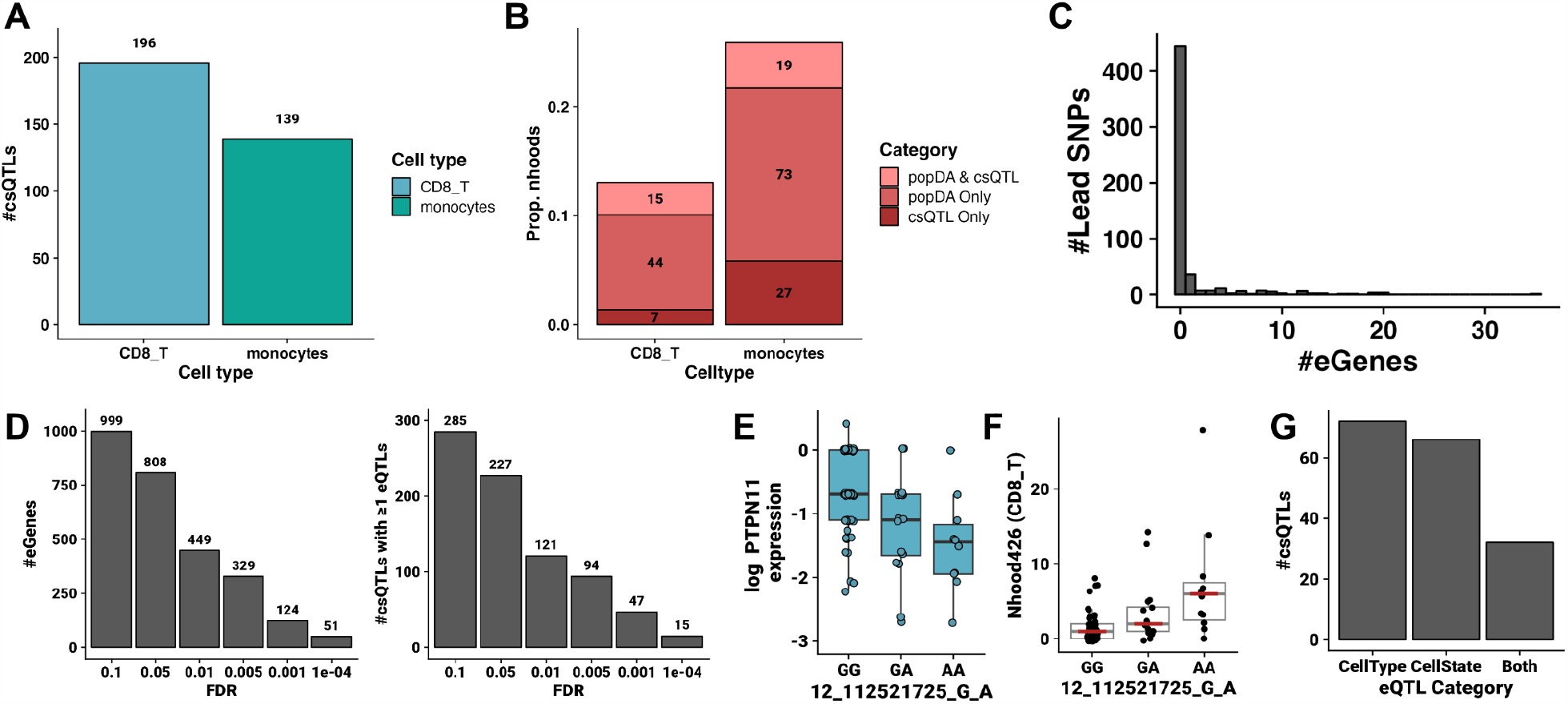
Cell state QTL identification across immune cells using Milo2.0. (A) csQTLs identified across peripheral immune cells from Randolph *et al*. Each bar represents the number of lead SNPs identified in each immune cell type (colour as in Figure 2A). (B) Proportions of neighbourhoods that are either csQTLs, popDA cell states or both, for each immune cell type. Numbers are computed relative to the number of neighbourhoods present in each category. (C) Distribution of eQTL Catalogue eGenes linked to csQTLs by colocalization analysis. (D) Cell state eQTL analysis identifies diminishing eGenes (left) and eQTLs (right) across FDR thresholds. (E) Neighbourhood log expression of *PTPN11* vs. SNP allele in neighbourhood 426 corresponding to a CD8+ T cell state. (F) Cell abundance corresponds to the same neighbourhood and SNP in (E). (E-F) Boxplots show the median +/-interquartile range (IQR) with whiskers extending 1.5x IQR. Individual data points n=89 samples are shown. (G) Numbers of csQTLs that colocalise with at least one eQTL in either cell states or cell types from the eQTL catalogue or both.

To address if csQTLs were driven by eQTLs in the same cell types, we intersected our results with eQTLs identified by Randolph *et al*. Across a range of spatial FDR thresholds, we found a single overlap with our csQTLs (1/335, 0.35% FDR 1x10^-8^), indicating that they are unlikely to be driven by eQTLs in the corresponding cell type cluster (Supplementary Figure 9A). We further explored this by performing statistical colocalization analysis [29] with immune cell eQTLs from the eQTL catalogue [30]. We identified a small number of csQTLs that could be explained by eQTLs in equivalent cell types (68/576, 11.8%, Figure 3C). These results suggest that either highly cell-state specific eQTLs, or other molecular mechanisms, explain our csQTLs. We reasoned that if heterogeneity of molecular states within a cell type (i.e. the existence of additional substructure) resulted in a lack of sensitivity to detect eQTLs, it would be interesting to explore whether cell state-specific eQTLs were present in genes proximal to each csQTL.

To this end, we identified all protein-coding genes within 1Mb of each lead csQTL SNP and explored the gene expression patterns across the relevant csQTL neighbourhood. Of the set of csQTL proximal genes, 1684 (37.1%) were expressed in at least one csQTL neighbourhood. Of these, we found (using a pseudo-bulk approach; Methods) that 449 (26.7%) had at least 1 eQTL (FDR 1%), corresponding to 21.0% (121/576) of our csQTL signals (Figure 3D, Supplementary Figure 9B). Moreover, in a colocalization analysis we found strong (PP > 0.75) evidence of shared csQTL and eQTL signals for 98/576 (17.0%) csQTLs (Supplementary Figure 9C-D). For example, 10 csQTLs corresponding to a single locus colocalize with an eQTL for the protein tyrosine phosphatase SHP2, encoded by *PTPN11* (Figure 3E, Supplementary Figure 10). The presence of the minor allele of these csQTLs increases the abundance of CD8+ T cell states with subsequently lower expression of *PTPN11* (Figure 3F, Supplementary Figure 11), demonstrating the complementary information gained from mapping both csQTLs and eQTLs in the same samples. In sum, either cell type or cell state eQTLs colocalize with 170/576 (29.5%) csQTLs (Figure 3G). These analyses indicate that the majority of csQTLs may not be driven by eQTLs in corresponding cell types or states, warranting the investigation of alternative molecular and cellular mechanisms in the future.

### Cell state QTLs link single-cell gene expression to human traits

Next, we used colocalization analysis to explore whether immune csQTLs were associated with GWAS summary statistics for human diseases and traits. Comparing our csQTLs to the summary statistics from the pan-UKBB [https://pan.ukbb.broadinstitute.org] and FinnGene studies [31] we found strong evidence (PP ≥ 0.75) that 261 of the csQTLs colocalized with 218 traits (Figure 4A, Supplementary Figures 12-13). Focussing on traits with immune system involvement, we identified csQTLs that could be linked to a range of traits and diseases (Figure 4B, Supplementary Figure 12D, 13D), including sepsis, immune cell abundance (eosinophils, neutrophils and lymphocytes), autoimmunity (juvenile arthritis, psoriatic arthritis, Sjögrens), allergies (eczema, allergic contact dermatitis) and susceptibility to a range of infectious diseases (e.g. tick-borne encephalitis, mumps). To link gene regulation, immune cell states and disease susceptibility we considered the set of csQTLs that were colocalized with both cell state eQTLs and human immune traits (Figure 4C; Supplementary Table 2). For example, genetic regulation of *SH2B3* expression in CD8 T cells, and the abundance of *SH2B3*+ CD8 T cells, is colocalized with lower risk of connective tissue disease (Figure 4D, Supplementary Figure 14A-C). *SH2B3* encodes Lnk, an adaptor protein critical for the negative regulation of cytokine and lymphocyte antigen receptor signalling [32]. Our results indicate that genetically reduced expression of SH2B3 may lower the threshold of activation of CD8+ T cells in response to cytokines, for example IL-15 [33]. Consequently, CD8 T cells that are both more abundant and more sensitive to cytokine cues may provide protection against hypothesised infectious triggers of connective tissue disease, such as COVID19 [34,35]. Similarly, we identified a CD8 T cell csQTL colocalized with lower *RDH14* expression, that encodes a retinol dehydrogenase, and that is also associated with an increased risk of mumps, a paediatric viral infection (Supplementary Figure 14D-F). In summary, our analyses demonstrate the added value of csQTLs for functionally interpreting autoimmune and infectious disease associated loci. More broadly, we present a workflow to identify cell state QTLs that can provide novel insights into the cellular genetic basis of human disease.

**Figure 4.**
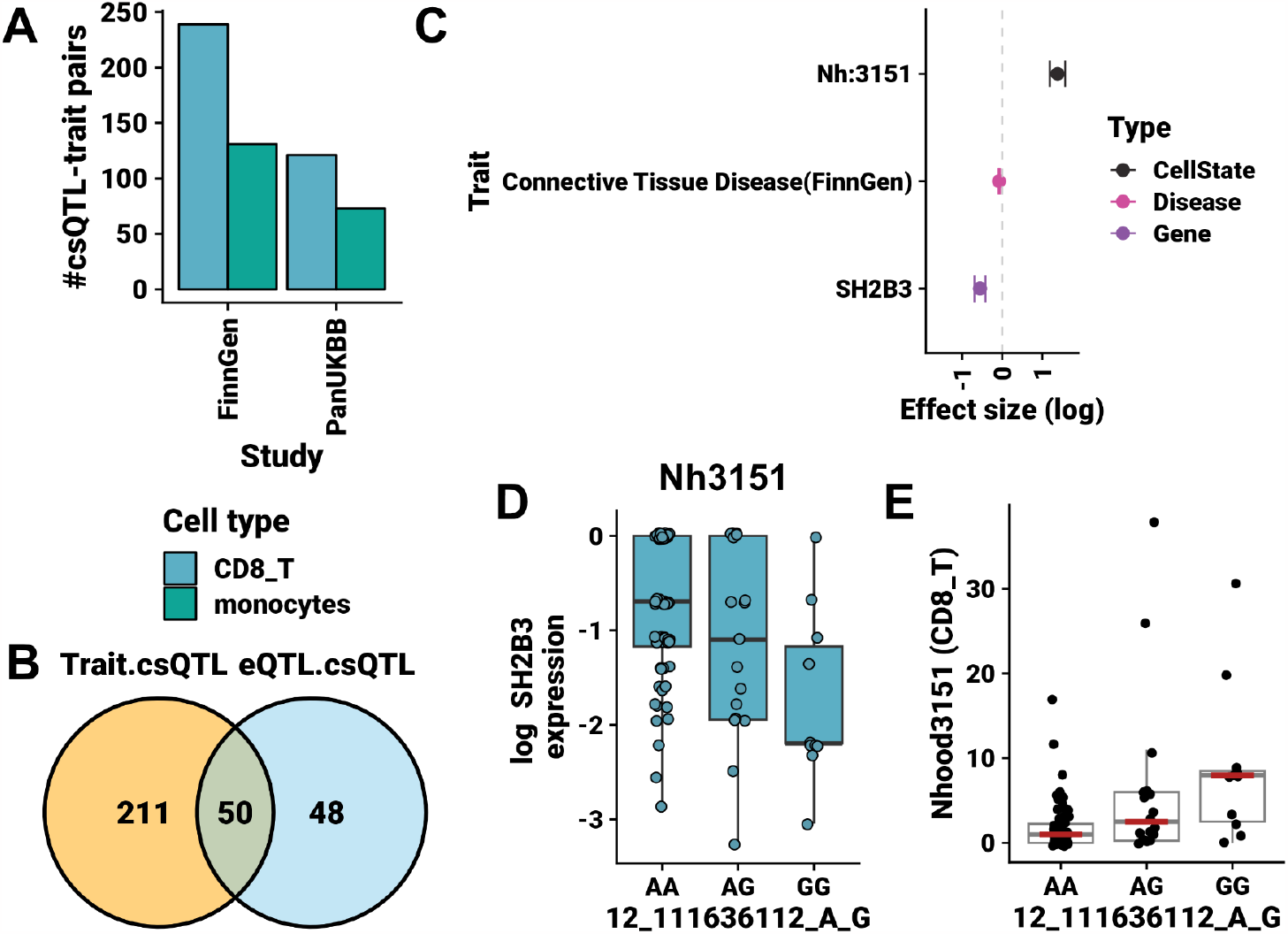
Immune cell state QTLs colocalize with human health traits. (A) Colocalization of csQTLs with human traits across the FinnGen and Pan-UKBB studies. Bars are coloured by the cell type to which the cell states map. (B) Venn diagram of the intersection of csQTLs colocalized with traits and/or eQTLs. (C) Effect size estimates used for colocalization analysis linking *SH2B3* expression, CD8 T cell state abundance (neighbourhood 3151) and connective tissue disease risk. Points are model estimates with 95% confidence intervals. (D) Boxplots denoting the expression of *SH2B3* across samples in neighbourhood 3151 for the csQTL denoted in (C). Points show the average expression of *SH2B3* across single cells from each individual (n=89). (E) Cell abundance for the equivalent neighbourhoods shown in (D). (D-E) Boxplots show the median +/-interquartile range with whiskers extending 1.5x IQR. Individual data points n=89 samples are shown.

## Discussion

We introduce Milo2.0, a significant advance on Milo for the identification of perturbed cell states from single-cell studies. Milo2.0 provides a scalable framework to integrate population-scale single cell data sets with statistical genetic analysis. Demonstrating its utility, we use Milo2.0 to identify both ancestry-DA immune cell states and cell state QTLs (Figures 2 & 3).

The identification of csQTLs has previously been limited to flow cytometry-based studies, which require the optimisation of multiplexed antibody panels to quantify cell types and states of interest [1,2,10,11]. These studies have provided strong evidence for the genetic regulation of cell type abundance, but have been limited to studying cell types based upon the expression of pre-defined surface antigens. More recently, population-scale single cell data sets have used annotated clusters to represent cell types and attempted to identify genetic regulation of their abundance [5,6]. As we have previously noted [20], this may be suboptimal for identifying factors driving cell state variation due to inherent limitations in cluster-based representation of cell states. We overcome the limitations of both study approaches by representing cell states as overlapping kNN-graph neighbourhoods and demonstrate the genetic regulation of cell state abundance at the high resolution afforded by scRNA-seq (Figure 3).

While we introduce algorithmic advances that increase the scalability of the Milo workflow to millions of single cells (Figure 1), the NB-GLMM introduces a new limitation. Namely, the REML projection matrix requires a series of dense matrix multiplications that limit the scalability of our implementation to data sets of N=1000 or fewer individuals (best case scaling is ∼O(n^2.4^) and worst case is O(n^3^). Overcoming this analytical bottleneck may be achieved by introducing advances in Variational Bayes to the NB-GLMM [19], leveraging graphical processing units (GPU), implementing auto-differentiation [36], or identifying a factorisation of the Negative Binomial likelihood or pseudo-likelihood akin to the Gaussian case [13]. Currently, the largest published single-cell population cohort study contains ∼1000 individuals [6], but many other studies are an order of magnitude smaller [7,8]. As cohort sizes expand we anticipate the need for faster algorithms and meta-analytical frameworks, particularly for federated analyses that are becoming common for human genetic analyses [37].

We have identified > 500 csQTLs in resting peripheral immune cells (Figure 3, Supplementary Table 1). Intuitively, we might expect these to be driven by genetic regulation of gene expression in the same cell states. While, we find ∼20% of csQTLs can be explained by eQTLs, including those that colocalise with human traits and diseases, our results indicate a more complex regulatory picture, which may include, for example, splicing QTLs [38]. Many of the “unexplained” csQTLs (211/576, 36.6%) in our study also colocalize with human traits (Figure 4B). This result might be due to genetic effects acting upstream of cell states, i.e. in their differentiation or maturation trajectories, suggesting genetic regulation of cell fate choices may be an unexplored aspect of genetic predisposition to disease. This is concordant with the emerging concept that response eQTLs may explain GWAS findings better than homeostatic eQTLs [39]. In summary, we have demonstrated the integration of differential abundance testing with statistical genetic analysis, and provide the framework to facilitate the identification of novel genetic regulation of cell states relevant to human health and disease.

## Methods

### Counting triangles as an approximation to neighbourhood distances

Milo seeks to identify a set of kNN graph neighbourhoods that are representative of cell states in a single-cell experiment. To achieve this, while dispensing with the need for computationally expensive distance calculations, we have implemented a graph-based approximation to identify these neighbourhoods. For a graph *G*(*V, E*) we randomly sample a proportion *P* of the vertices such that the sampled vertices *P* _*v*_ ⊂ *V*. For each *v* ∈ {1, 2, …, *P* _*v*_ } we create the induced subgraph *G*_*v*_ (*V*_*v*_, *E*_*v*_) comprised of all vertices that are connected by an edge to *v* in the graph *G*. Subsequently, we count the number of 3-step cyclic random walks (triangles) on *G*_*v*_ for each vertex using the algorithm in [22], as implemented in the function *count_triangles* from the R package *igraph* [40]. The refined neighbourhood index is defined as the vertex *V*_*v*_ which has the highest number of triangles in *G*_*v*_.

### Number of shared cells for spatial FDR correction

Milo uses a false discovery rate computed as the weighted Benjamini-Hochberg adjusted p-value. Weights are computed as the reciprocal kth nearest neighbour distance to the neighbourhood index cell. The justification for this approach follows the logic in [41], such that the weight accounts for the overlap of neighbourhoods in the N-dimensional reduced dimension space (e.g. principal component space). To circumvent the need for distance calculations we introduce an alternative weighting scheme using the number of overlapping cells between each neighbourhood. Specifically, for each refined neighbourhood we count the number of overlapping cells with all other neighbourhoods and compute the adjustment weight as the reciprocal of this sum.

### Negative binomial GLMM

We extend the Milo DA testing framework by enabling a negative binomial-GLMM (NB-GLMM). This allows additional flexibility in experimental designs by incorporating random effect variables that account for dependence between observations that might otherwise lead to inflated type I errors. To define an NB-GLMM we require several components: (1) a linear predictor *η* = *Xβ* + *Zb*, where *X* is the fixed effect design matrix, mapped onto a set of weights (*β*) and *Z* is the random effect design matrix with weights, *b*, (2) a sampling distribution for the conditional observations, *y*|*b*∼*NegativeBinomial*(*μ, ϕ*), (3) a sampling distribution for the random effects, *b*∼*Normal*(0, *G*), and (4) a link function, *g*(*μ*|*b*) = *η* = *log*(*μ*|*b*) with inverse function, *h*(*η*) = *e* ^*η*^, which maps the model space (*η*) onto the data space (μ). To normalise our NB-GLMM for variation in the number of cells sampled per observation we include an offset term as we do for the NB-GLM; by default this is computed from the trimmed mean of M-values (TMM) [42] such that *μ* = *e*^(*log*(*O*)+ *Xβ* + *Zb*)^, where *O* is the offset (TMM or otherwise). Further details on the negative binomial distribution can be found in the Supplementary Note.

### Pseudo-likelihood GLMM

Solving linear mixed models requires marginalising over the random effects, which is typically a high-dimensional and intractable integral. To overcome this issue, the pseudo-likelihood was proposed [24]. Pseudo-likelihood approximates the high-dimensional integral in a similar vein to Laplace approximation; both are motivated by a Taylor expansion. For the pseudo-likelihood, the expansion is on the inverse link function, *h*(*η*), evaluated at the predictor 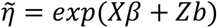 as described in [24], which is then used to compute a working pseudo-variable *y*^∗^(see Supplementary note for details). This yields a term for the expected value 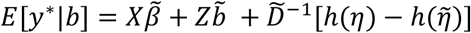 and the variance 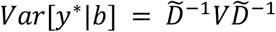, where *y*^∗^|*b*∼*Normal*(μ, *V*^∗^(*σ*)). We then proceed with fixed and random effect parameter estimation using an adaptation of Henderson’s mixed model equations (MME) for the pseudolikelihood:

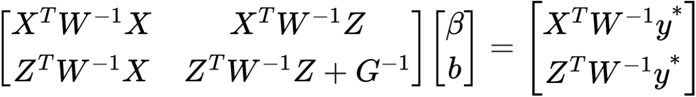

Where *W* = *D* ^− 1^*V* _μ_ *D*^− 1^and 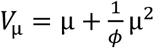 which is the negative binomial variance. We also define the pseudo-variance:

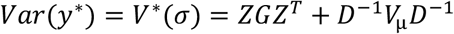

And the pseudo-log-likelihood:

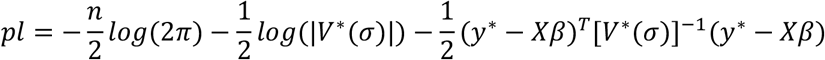

### Fisher scoring

For variance parameter estimation we use the restricted maximum likelihood (REML), which maximises the likelihood after accounting for the fixed and random effect weight parameter estimation, and for which the pseudo-log-likelihood is:

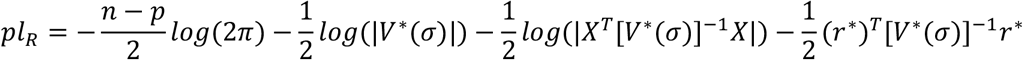

Estimation iteratively updates the current estimate based on the gradient (score) and the Hessian (information) matrix:

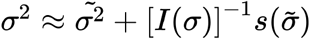

We define the score vector for the i^th^ variance component, *s*_*i*_(*σ*):

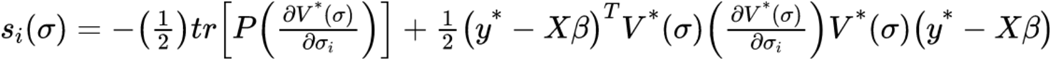

Further, the ij^th^ component of the information matrix is defined as:

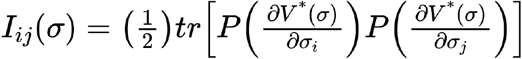

Where *P* = [*V*^∗^(*σ*)]^− 1^ *−* [*V*^∗^(*σ*)]^− 1^*X*(*X*^*T*^[*V*^∗^(*σ*)]^− 1^*X*)^− 1^*X*^*T*^ [*V*^∗^(*σ*)]^− 1^, is the REML projection matrix and 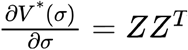 is the partial derivative of the pseudo-variance with respect to each variance component. Estimation of variance components then proceeds iteratively with initial estimates of *Θ*_0_ = [*β b σ*^2^], followed by oscillating estimation of [*β b*] by the MMEs as described above, and Fisher scoring to update *σ*^2^. The convergence criteria at the *i*th iteration is 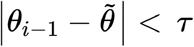, where 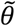 is the current estimate and *τ* = 10^− 5^ by default.

### Haseman-Elston regression and non-negative least squares

Fisher scoring is a quasi-Newton approach, and thus can converge quickly to a (local) optimum. However, when *c* → *N*, the information matrix will also tend to zero, with an undefined inverse. In practice, this occurs when a predefined *n x n* covariance matrix is provided, such as a genetic relationship matrix. To address this, we re-frame the variance parameter estimation in terms of a least squares regression. This approach, called Haseman-Elston regression, was proposed for sib-difference regression [43,44]. Here we use it to decompose the pseudo-covariance into *c* + 1 linear variables, i.e. variance components. First, we vectorise the columns of the *n x n* REML pseudo-covariance matrix *Var*[*Y*^∗^] = *P*[*y*^∗^*y*^∗*T*^]*P*, which is equivalent to stacking the n columns into a single n^2^ vector. In practice, *Var*[*Y*^∗^] is symmetric, so we only need to stack the n(n-1)/2 elements including the diagonal elements. Accordingly, we also stack the columns of *ZZ*^*T*^ for each random effect *j* ∈ {1, 2, …, *c*}, to define our new regression problem:

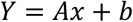

where *Y* = *vec*(*P*[*y*^∗^*y*^∗*T*^]*P*)

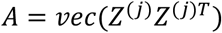

The solution to the linear system of equations can be solved via least-squares, where the vector of 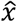 are the estimated variance components. For the case with a genetic relationship matrix, or other user-defined covariance matrix *K*, we vectorise the columns in the same manner as for *P*[*y*^∗^*y*^∗*T*^]*P*.

Variance parameters are constrained to the positive real numbers, *σ*^2^ ∈ [0, ∞]. During optimization, variance estimates may stray outside the lower boundary, particularly for small sample sizes and for variances close to zero. Different approaches have been taken to handle this boundary problem, including manual constraint of parameters. However, this may lead to convergence issues as there is no guarantee that this bounding is optimal. Therefore, we adopt a non-negative least squares (NNLS) solver, which uses active and passive sets to optimise the non-constrained parameters [45]. Convergence is checked by conditions on the Lagrangian multipliers such that *ω* _*j*_ = 0 *j* ∈ *ϒ* and *ω*_*j*_ ≤ 0 *j* ∈ *Λ*, where *ϒ* corresponds to the unconstrained parameter set and Λ is the set of constrained parameters [45]. In our implementation, if any variance parameter estimates are < 0 the model will automatically switch to HE-NNLS to prevent failure. We find that Fisher scoring, HE and HE-NNLS give equivalent results on simulations (Supplementary Figure 15).

### Dispersion parameter estimation

We adopt a 2-stage approach to dispersion parameter estimation. In the first off-line stage we compute neighbourhood-specific dispersion values using an approximate profile likelihood approach, as implemented in *edgeR*. This maintains consistency with the NB-GLM for fixed effect models. For the NB-GLMM, and to avoid the neighbourhood count variances being absorbed by the initial dispersion estimate (which is agnostic to the model design), we re-estimate each neighbourhood dispersion parameter separately using a golden section search, with an upper bound set by the initial *edgeR* estimate; the lower bound is set to 50% of the initial estimate. The second on-line stage then uses methods of moments to calculate an updated dispersion parameter at each iteration of the NB-GLMM solver. Methods of moments use the relationships between parameters with known estimators to calculate a value for a parameter without a known estimator. In our NB-GLMM we use the fact that the sample variance 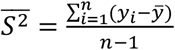 is a maximum likelihood estimate for the variance, and that 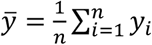 is a maximum likelihood estimate for the mean. Using the relation of the NB-GLMM pseudo-variance *Var*[*Y*^∗^] = *D*^− 1^ *V* _μ_ *D*^− 1^ + *ZGZ*^*T*^ where *D*^− 1^ = 1/μ is an *n x n* diagonal matrix and 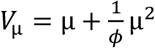 is the negative binomial variance, and substituting 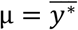 and 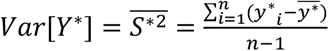, then rearranging for *ϕ* we get:

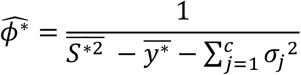

### Inference and hypothesis testing

The primary goal of Milo is to identify DA neighbourhoods. In fixed-effect only mode this uses the quasi-likelihood F test implemented in *edgeR* [46,47]. Our NB-GLMM uses a pseudo-likelihood, which does not permit the use of likelihood ratio tests. Therefore, we perform inference on our fixed effect variables using Wald-type statistics, described below. We first compute a test statistic 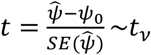, where 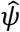 is our fixed effect parameter estimate, *ψ*_0_ is the null hypothesis value where generally *ψ*_0_ = 0. The standard error of the parameter estimate, 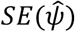, is computed directly from Hessian matrix where 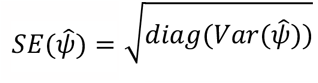. The t-distribution has *v* degrees of freedom, which for a fixed effect-only model with *n* observations and *p* parameters is *n − p*. However, for a mixed effects model the degrees of freedom need to be approximated. We use the Satterthwaite approximation, which has been shown to have good small-sample behaviour and equivalent performance to the Kenward-Roger approximation [48]. Specifically, we compute:

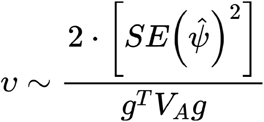

Where, 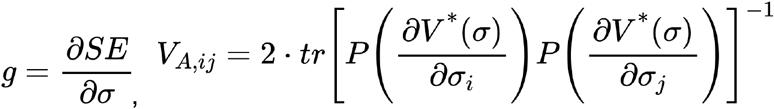 and *P* is the REML projection matrix defined above. We then compute *p*_*nh*_ = *P*(|*t*| ≥ *t*_*v*_) = 2 · [1 *− T*(*t*; *v*)], where *T*(⋅ ; *v*) is the distribution function for the *t*_*v*_ distribution.

### Simulations and benchmarking

Single cell data sets were simulated as described previously using *dyntoy* [49]. To generate 1 million single cells, we simulated 5 random datasets of 200,000 cells each using *dyntoy* and merged the data. Neighbourhood counts from unrelated individuals were sampled from a negative binomial distribution in R (*rnbinom* function) parameterised by the mean and dispersion: ⌊ *C*_*i*_ ∼*NB* (*μ*_*i*,_ *ϕ)*⌋ for *i* ∈ {1,…,*N* } observations.

Where, *μ*_*i*,_ =exp (*x*_*i*_ *β* + *z*_*i*_*b*)

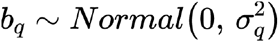 for *q ∈* {*1*,…,*q*} random effect variables with the design matrix

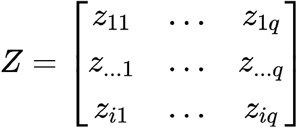

*β* = [*β*_1_,…, *β*_*k*_] for {*1*,…,*k*} fixed effect variables with the design matrix

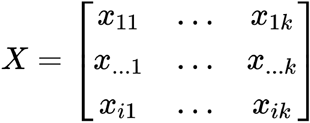

The floor operator ⌊ … ⌋ was used to retain the integer nature of neighbourhood counts. Weights for 2 fixed effect variables were in the intervals *β*_1_=[-1, 1.2] in steps of 0.4 and *β*_2_=[0.1, 1] in steps of 0.1 as ground truth values. Similarly, ground truth random effect variances (σ^2^_1_, σ^2^_2_) were selected for 2 random effect variables in the interval [0.5, 1.5] in 0.1 step sizes. N=150 individuals were simulated for each combination of ground truth values to create 90 simulated data sets.

For each dataset we ran Milo2.0 alongside *glmmTMB* [36] and *glmmPQL* from the *lme4* R package [50]. For Milo2.0 we used Fisher scoring as the solver and provided an estimate of the dispersion computed using methods of moments (see Supplementary Note); *glmmTMB* and *glmmPQL* compute estimates of ϕ internally. Both *glmmTMB* and *glmmPQL* used a log-link model with the *nbinom2* parameterisation, which most closely matches that used in Milo2.0. All other model options were set as default. Model parameter estimates and p-values were extracted from the respective result objects. Parameter estimates were compared to ground truth using the absolute deviation and p-values were compared across models for each fixed effect parameter.

To simulate related individuals, we constructed P random pedigrees containing between 1 and 3 generations with defined genetic relationships, i.e. sib-sib=0.5, sib-parents=0.5, parent-parent=0, sib-grandparent=0.25. Each pedigree contained at least 5 individuals. For each individual in each family, we simulated counts for a single neighbourhood using the following parameters: β=[0.3, 0.7] for 2 fixed effects; 1 binary and 1 continuous where the continuous fixed effect variable was simulated using *rnorm* with mean=0 and sd=1, σ^2^=[0.4, 0.4] for 2 random effect variables with 5 and 4 levels, respectively. Pedigrees were assigned a unique pedigree ID as an unordered categorical variable. To simulate the effect of genetic relationships on the counts distribution we projected the vector of simulated counts *Y* onto the Cholesky decomposition of the total variance matrix to obtain *Y*^*g*^:

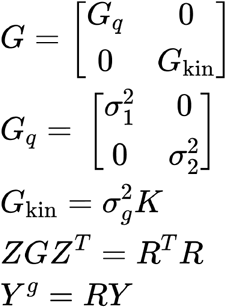

Here *G* is the full covariance matrix across all individuals and families for 2 random and 1 genetic effect. *K* is the N x N kinship matrix of known genetic relationships from the simulated pedigrees, which is computed by concatenating together individual pedigree kinships, assuming between pedigree relationships = 0. *R*^*T*^*R* is the upper-triangular Cholesky decomposition computed in R using the *chol* function. *Y*^*g*^ was used as input to *Milo2*.*0, glmmTMB* or *glmmPQL* using the family ID as the random effect. Additionally, the kinship matrix *K* was input directly into Milo2.0 as the sole random effect.

### Data set processing

Single cell data from Randolph *et al*. was downloaded from GEO (accession GSE162632) as count matrices. Size factors were estimated using *scran* [51], highly variable genes (HVGs) were computed and used as input to principal components analysis (PCA) prior to batch integration with *fastMNN* [52] prior to Milo analysis. Alternatively, single cells from mock and IAV infected samples were split after size factor estimation and normalisation, but prior to HVG, PCA and batch integration steps. These latter steps were performed separately on the split data prior to separate Milo analyses.

Data from Stephenson *et al*. were downloaded from the COVID19 Cell Atlas portal (https://www.covid19cellatlas.org/index.patient.html). Single-cells were subset to T cell annotations: *CD4*.*Activated, CD4*.*CM, CD4*.*EM, CD4*.*Naive, CD4*.*Prolif, CD4*.*Tfh, CD4*.*Th1, CD4*.*Th2, CD4*.*17, Treg, CD8*.*EM, CD8*.*Naive, CD8*.*Prolif, CD8*.*TE, gdT, MAIT* and *NKT*. Batch correction was performed using fastMNN [52] (k=60), prior to Milo analysis.

### Population differential abundance testing analysis

We identified population differential abundant (popDA) cell states using Milo2.0. We computed the k-nearest neighbour graph across the respective data sets: Randolph mock-infected (k=50, dimensions=50), Randolph IAV-infected (k=50, dimensions=50), Stephenson COVID19 T cells (k=60, dimensions=30). Each graph was used as input, along with the respective batch-corrected reduced dimensions, computed using fastMNN [52], to generate neighbourhoods that captured cell state variation and allowed for sufficient counts across neighbourhoods and study samples. All Milo workflow parameters are recorded in Supplementary Table 3.

### Cell state quantitative trait locus analysis

We performed genome-wide cell state QTL discovery using Milo2.0. Post-QC imputed genotype data were acquired from Randolph *et al* [9]. Common SNPs (sample MAF ≥ 15%) were selected for analysis and subsetted to SNPs that did not segregate between populations, i.e. contained non-zero heterozygous or minor allele homozygous allele counts, retaining m=4,638,095 SNPs for analysis. Neighbourhood count distributions were fit across all non-duplicated individuals for each Milo neighbourhood in the mock-infected control samples (N=89). The Milo2.0 NB-GLMM was fit for each neighbourhood separately, adjusting for genetic relationships two-fold: (1) non-independence was accounted for as a random effect using genetic relationship matrices (GRM) computed from all SNPs with a leave-one chromosome out (LOCO) approach to exclude the chromosome for which the test SNP was assigned, (2) population structure confounding with mean neighbourhood count was adjusted using the self-reported genetic ancestry variable from Randolph *et al*. as a fixed effect. The spatial false discovery rate was computed separately for each SNP accounting for multiple testing across neighbourhoods. Across all SNPs tested we defined genome-wide significance (spatial FDR ≤ 1x10^-8^) as evidence of a cell-state QTL.

For each chromosome, we first considered all SNPs that reached genome-wide significance (spatial FDR ≤ 1x10^-8^) across all neighbourhoods. For each neighbourhood, loci with multiple associated SNPs were clumped together if they were within +/-500kb of the most strongly associated SNP. This aggregation process collapsed all 32,574 initial csQTLs down to 576 final csQTLs.

### Expression quantitative trait locus analysis

We identified eQTLs in the Randolph *et al*. neighbourhoods using a pseudo bulk approach. First, we identified the protein-coding genes with start or end coordinates within 1Mb of the lead SNP for each csQTL. We extracted the single cell gene count matrix for cells assigned to each Milo2.0 neighbourhood in the csQTL. To compute the pseudo bulk expression we aggregated the gene counts for each cell and gene within the respective 1Mb interval for each experimental sample, i.e. if A is the binary cell X sample matrix and B is the gene X cell counts matrix then the pseudo bulk counts matrix C is *C* = *A*^*T*^*B*. We then normalised the per gene expression profiles by dividing the pseudo bulk gene counts by the number of cells from each sample and performed a log transform with a +1 pseudocount added to both the numerator and denominator.

Normalised gene expression for each locus were used as input to *Matrix eQTL* testing [53] for all SNPs within 1Mb of the csQTL and 1Mb of the most distal genes within this interval. SNPs were removed with MAF < 15% in line with the csQTL testing approach described above. Each gene was tested for association with the SNPs in *cis*, bounded by the furthest SNPs from either the gene body or corresponding csQTL. The Matrix eQTL regression model was adjusted for ancestry, expressed as the proportion of estimated African ancestry from Randolph *et al*. [9]. An eQTL was defined as a statistically significant association (FDR 1%) between a SNP and gene.

### Colocalization analysis

Colocalization between csQTLs and either eQTLs or summary statistics from genome-wide association analyses was performed using the R package *coloc* [29]. Default prior probabilities were used (p1=10^-4^, p2=10^-4^, p12=10^-5^) as well as summary statistics from both csQTLs and the trait/eQTL to be colocalised. SNPs were selected within +/-1Mb of the lead csQTL SNP for both csQTL signal and trait, and the intersection of SNPs was analysed. Colocalization analysis was performed on loci for which an eQTL (FDR 1%) or trait association was evident (p≤10^-5^). Pan-UKBB SNPs are in GRCh37 coordinates, while csQTL, eQTL and FinnGen study SNP coordinates are all mapped to GRCh38. Therefore, we used the liftOver tool [https://genome.ucsc.edu/cgi-bin/hgLiftOver] to assign Pan-UKBB SNPs a corresponding coordinate on the GRCh38 human genome build. Allele harmonisation was carried out to check for concordance of reference and alternate alleles and, where necessary, swap alleles across strands to achieve harmonisation. Input summary statistics were the (log-scale) effect estimates, standard errors and, for quantitative traits, the trait standard deviation. If no trait variances or standard deviations were available we included the alternate allele frequency and sample. In the absence of regression estimate standard errors, these were estimated from the SNP p-values and sample size assuming they were drawn from a standard normal distribution. In all colocalization analyses we make the single causal variant assumption. Colocalized trait-pairs were defined by a posterior probability for hypothesis 4 ≥ 0.75, where each hypothesis is defined: H_0_ - neither trait 1 nor trait 2 show association signal, H_1_ - only trait 1 (csQTL) is associated, H_2_ - only trait 2 is associated, H_3_ - trait 1 and trait 2 are associated but with different SNPs, H_4_ - trait 1 and trait 2 associations are colocalised, i.e. explained by the same genetic signal.

## Code and data availability

Analytical code used in this study are included at: https://github.com/MorganResearchLab/Milo2.0Analysis_2023. All Milo software is hosted at https://github.com/MarioniLab/miloR and is available through Bioconductor. Summary statistics for csQTLs are available via Zenodo: 10.5281/zenodo.10075946.

## Acknowledgements

The authors thank S. Ghazanfar, K. Hua, E. Dann and other members of the Marioni lab for discussions regarding statistical and computational aspects of the project.

J.C.M. acknowledges core funding from EMBL which supported AK and core support from Cancer Research UK (C9545/A29580) which supported MDM. This work was supported by the Human Biomolecular Atlas Project (NIH 1OT2OD026673-01) to J.C.M.

