## Supplemental Note for "Milo2.0 unlocks population genetic analyses of cell state abundance using a count-based mixed model"

Notation

For clarity, we use the following notation to distinguish scalars, vectors and matrices. Scalars are generally denoted with lower case letters, such as ‘a’ or ‘c’, vectors are denoted by lower case bold letters or greek letter, e.g. the row vector $\mathbf{v} = [0 \quad 1 \quad 0]$ or $\beta = [\beta_1 \quad \beta_2 \quad \beta_3]$.

Matrices will be denoted by upper case bold letters, i.e. $\mathbf{X} = \begin{bmatrix} 1 & 0 \\ 0 & 1 \end{bmatrix}$.

Generalised linear (mixed) models for the negative binomial distribution

Using the Nelder and Wedderburn definition of a GLM [1], the negative binomial likelihood can be expressed in terms of an exponential family distribution when the dispersion parameter, $\phi$, is assumed known. Hence, an exponential family distribution likelihood function $f(y|\theta)$ can be expressed in the form:

$$f(y\,|\,\theta) = \exp\left[\frac{t(y)\theta - b(\theta)}{a(\gamma)} + c(y,\,\gamma)\right] \quad (1)$$

Where $t(y)$ is the sufficient statistic, $\theta$ is the canonical parameter, $b(\theta)$ is the cumulant generating function in terms of the canonical parameter, $c(y,\,\gamma)$ is a re-scaled non-negative function of $y$, and $a(\gamma)$ is a scale parameter for the relevant distribution.

The negative binomial likelihood function $L(\theta; y)$ for $\theta = [p,\,\phi]$ is:

$$L(p,\,\phi; y) = \binom{y + \phi - 1}{y}(1-p)^y p^{\phi} \qquad (2)$$

Therefore, we can write (2) in terms of (1), assuming $\phi$ is known:

$$L(p,\,\phi; y) = \exp\left[\underbrace{y\log\left[1-p\right]}_{\mathrm{t(y)}\theta} + \overbrace{\phi\log\left[p\right]}^{\mathrm{b}(\theta)} + \underbrace{\left[\sum_{i=1}^{k} y_i\right]^{-1}\log\left[\frac{\Gamma(y)}{\Gamma(\phi)}\right]}_{\mathrm{c(y,}\gamma)}\right] \text{where } k = \max(y_i) \quad (3)$$

Hence, $t(y) = y$, $\theta = log(1 - p)$, $a(\gamma) = 1$ and $b(\theta) =- \phi log(p)$. The GLMM is framed in terms of the conditional expectation and variance $E(y\,|\,b) = \mu\,|\,b$ and $var(y\,|\,b) = \mathbf{V}_{\mu}^{1/2}\mathbf{A}\mathbf{V}_{\mu}^{1/2}$, respectively. Here, $\mathbf{V}_{\mu}^{1/2} = diag\left[\sqrt{\frac{\partial^2 b(\theta)}{\partial\theta^2}}\right]$ and $\mathbf{A} = diag\left[\frac{1}{a(\gamma)}\right]$, which for the negative binomial parameterisation here leads to $\mathbf{A} = \mathbf{I}$ and therefore $var(y\,|\,b) = \mathbf{V}_{\mu}^{1/2}\mathbf{I}\mathbf{V}_{\mu}^{1/2} = \mathbf{V}_{\mu}$.

To find the expression for $\mathbf{V}_\mu$ we need to find the variance of the negative binomial in terms of μ and the canonical parameter θ.

For the random variable $Y \sim \text{NegativeBinomial}(p, \phi)$, $\mu = E[Y] = \dfrac{\partial b(\theta)}{\partial \theta}$ and $Var[Y] = \gamma \dfrac{\partial^2 b(\theta)}{\partial \theta^2}$. Recalling the term for $b(\theta)$ from (3) and setting $e^\theta = 1 - p$ we have $b(\theta) = -\phi \log\left[1 - e^\theta\right]$.

Let $u = 1 - e^\theta$, $v = \phi \log u$, then using the chain rule from calculus we find:

$$E[Y] = \mu = \frac{\partial b(\theta)}{\partial \theta} = \left(\frac{dv}{du}\right)\left(\frac{du}{d\theta}\right) = \left[-e^\theta \cdot \frac{-\phi}{1 - e^\theta}\right] = \frac{\phi e^\theta}{1 - e^\theta} \quad (4)$$

Substituting $e^\theta = 1 - p$ into (4) :

$$E[Y] = \mu = \frac{\partial b(\theta)}{\partial \theta} = \frac{\phi(1-p)}{p} \quad (5)$$

Continuing, we find an expression for $Var[Y]$ using (5):

$$Var[Y] = \gamma \frac{d^2 b(\theta)}{d\theta^2} = \gamma \frac{d(db(\theta)/d(\theta))}{d(\theta)} = \gamma \frac{d\mu}{d\theta} = \gamma \frac{d\frac{\phi[1-p]}{p}}{d\theta} = \gamma \frac{d\frac{\phi e^\theta}{1-e^\theta}}{d\theta}$$

Using the quotient rule: $\dfrac{df}{dt} = \dfrac{v\frac{du}{dt} - u\frac{dv}{dt}}{v^2}$

Let $u = \phi e^\theta and\, v = 1 - e^\theta$, with $\dfrac{du}{d\theta} = \phi e^\theta$ and $\dfrac{dv}{d\theta} = -e^\theta$

$$Var[Y] = \gamma \frac{d\frac{\phi e^\theta}{1-e^\theta}}{d\theta} = \gamma \frac{\left(1 - e^\theta\right) \cdot \phi e^\theta - \left(\phi e^\theta \cdot -e^\theta\right)}{\left(1 - e^\theta\right)^2}$$
$$= \gamma \frac{\phi e^\theta - \phi\left(e^\theta\right)^2 + \phi\left(e^\theta\right)^2}{\left(1 - e^\theta\right)^2} = \gamma \frac{\phi e^\theta}{\left(1 - e^\theta\right)^2} \quad (6)$$

Substituting $e^\theta = 1 - p$ into (6):

$$Var[Y] = \gamma \frac{d\frac{\phi e^\theta}{1-e^\theta}}{d\theta} = \gamma \frac{\phi[1-p]}{p^2} \quad (7)$$

For the GL(M)M We wish to express (4) and (7) in terms of μ:

$$\mu = \frac{\phi(1-p)}{p}$$
$$1 - p = \frac{\mu}{\mu+\phi}$$
$$p = \frac{\phi}{\mu+\phi}$$
$$\text{Substituting } 1 - p = \frac{\mu}{\mu+\phi} \text{into (7) :}$$

$$Var[Y] = \frac{\phi\left(\frac{\mu}{\mu+\phi}\right)}{\left(\frac{\phi}{\mu+\phi}\right)^2} = \frac{\frac{\mu\phi}{\mu+\phi}}{\frac{\phi^2}{(\mu+\phi)^2}} = \frac{\phi\mu}{\mu+\phi}\frac{(\mu+\phi)^2}{\phi^2} = \frac{\mu^2+\phi\mu}{\phi} = \frac{\mu^2}{\phi} + \mu \quad (8)$$

Where (8) is the familiar term for the negative binomial variance involving the quadratic mean scaled by the dispersion parameter $\phi$ (or the reciprocal in this case). Hence as $\lim_{\frac{1}{\phi}\to 0}$ then (8) collapses to the Poisson variance.

Pseudo-likelihood and pseudovariable definition

We make use of the pseudo-likelihood defined by Wolfinger and O'Connel [2], that transforms the observed data into a pseudovariable, which is approximately normally distributed. This facilitates the use of an adaptation of Hendersons' mixed model equations (MME) for parameter estimation [2,3]. For clarity, several terms from the methods are repeated here:

- The linear predictor: $\eta = \mathbf{X}\beta + \mathbf{Z}\mathbf{b}$
- The conditional sampling distribution of the observations: $\mathbf{y} \mid b \sim \text{NegativeBinomial}(\mu, \phi)$
- The sampling distribution of the random effect variables: $\mathbf{b} \sim Normal(0,\, \mathbf{G})$
- A link function: $g(\mu|b) = \eta = log(\mu|b)$
- An inverse link function: $h(\eta) = e^{\eta}$

We start with the partial derivative of the inverse link function with respect to the linear predictor, evaluated at the current estimate $\tilde{\eta}$:

$$\frac{\partial h(\tilde{\eta})}{\partial \tilde{\eta}} = h'(\tilde{\eta})$$
$$\tilde{\mathbf{D}} = diag\left[h'(\tilde{\eta})\right] = diag\left[e^{\tilde{\eta}}\right]$$

We write the first-order Taylor series expansion around the inverse link function at this estimate:

$$h(\eta) \approx h(\tilde{\eta}) + \tilde{\mathbf{D}}(\eta - \tilde{\eta})$$

Substituting the inverse link function in terms of the model parameter estimates we get:

$$\mathbf{X}\beta + \mathbf{Z}b \approx \mathbf{X}\tilde{\beta} + \mathbf{Z}\tilde{b} + \tilde{\mathbf{D}}^{-1}(h(\eta) - h(\tilde{\eta}))$$

We now define the pseudo-variable:

$$\mathbf{y}^{\cdot} = \eta + \tilde{\mathbf{D}}^{-1}[\tilde{\mathbf{y}} - h(\tilde{\eta})]$$

Recalling that the target of a GLMM is $E[y|b]$ and requires $Var[y|b]$ we define the equivalent expected and variance expression in terms of the pseudo-variable $y^*$:

$$E(\mathbf{y}^{\cdot} \mid b) = \mathbf{X}\beta + \mathbf{Z}\tilde{b} + \tilde{\mathbf{D}}^{-\tilde{1}}[h(\eta) - h(\tilde{\eta})]$$

$$Var(\mathbf{y}^{\cdot} \mid b) = \tilde{\mathbf{D}}^{-1}\mathbf{V}_{\mu}\tilde{\mathbf{D}}^{-1}$$

<u>Covariance matrix inversion</u>

Multiple steps during the GLMM parameter estimation require the inversion of a dense *n x n* covariance matrix, e.g. REML projection matrix, MME solving. The upper-bound on the number of operations is $O(n^3)$, hence mixed effect models have a reputation for poor scalability. While advances have been made for LMMs [4], to our knowledge equivalent refactorisations have not yet been developed for NB-GLMM or pseudo-likelihood approaches. In Milo 2.0, we use knowledge of the structure of the variance terms to implement faster matrix inversion functions.

Beginning with a result from [5] on the inverse of a sum of matrices:

$$\left(\mathbf{A} + \mathbf{U}\mathbf{B}\mathbf{U}^{\mathbf{T}}\right)^{-1} = \mathbf{A}^{-1} - \mathbf{A}^{-1}\mathbf{U}\mathbf{B}\left(\mathbf{I} + \mathbf{U}^{\mathbf{T}}\mathbf{A}^{-1}\mathbf{U}\mathbf{B}\right)^{-1}\mathbf{U}^{\mathbf{T}}\mathbf{A}^{-1}.$$

We note the similarity to the pseudo variance $\mathbf{V}^{\cdot}(\sigma) = \mathbf{W} + \mathbf{Z}\mathbf{G}\mathbf{Z}^{\mathbf{T}}$ where $\mathbf{W} = \mathbf{D}^{-1}\mathbf{V}_{\mu}\mathbf{D}^{-1}$. Hence, we can rewrite $\mathbf{V}^{\cdot}(\sigma)^{-1}$ in terms of $\mathbf{A} = \mathbf{W},\ \mathbf{U} = \mathbf{Z},\ \mathbf{B} = \mathbf{G},\ \mathbf{U}^{\mathbf{T}} = \mathbf{Z}^{\mathbf{T}}$:

$\mathbf{V}^{\cdot}(\sigma)^{-1} = \mathbf{W}^{-1} - \mathbf{W}^{-1}\mathbf{Z}\mathbf{G}\left(\mathbf{I} + \mathbf{Z}^{\mathbf{T}}\mathbf{W}^{-1}\mathbf{Z}\mathbf{G}\right)^{-1}\mathbf{Z}^{\mathbf{T}}\mathbf{W}^{-1}$, which involves computing 2 inverse terms $\mathbf{W}^{-1}$ and $\left(\mathbf{I} + \mathbf{Z}^{\mathbf{T}}\mathbf{W}^{-1}\mathbf{Z}\mathbf{G}\right)^{-1}$. We can trivialise the former by noting that $\mathbf{D}^{-1}$ and $\mathbf{V}_{\mu}$ are both diagonal matrices, hence their inverses can be computed in linear time (assuming all elements are non-zero). Indeed, we can simplify them further by evaluating directly:

$$\mathbf{D}^{-1}\mathbf{V}_{\mu}\mathbf{D}^{-1} = \begin{bmatrix} \frac{1}{\mu_1} & 0 & 0 & 0 \\ 0 & \frac{1}{\mu_2} & 0 & 0 \\ 0 & 0 & \ddots & 0 \\ 0 & 0 & 0 & \frac{1}{\mu_n} \end{bmatrix} \begin{bmatrix} \frac{\mu_1^2}{\phi} + \mu_1 & 0 & 0 & 0 \\ 0 & \frac{\mu_2^2}{\phi} + \mu_2 & 0 & 0 \\ 0 & 0 & \ddots & 0 \\ 0 & 0 & 0 & \frac{\mu_n^2}{\phi} + \mu_n \end{bmatrix} \begin{bmatrix} \frac{1}{\mu_1} & 0 & 0 & 0 \\ 0 & \frac{1}{\mu_2} & 0 & 0 \\ 0 & 0 & \ddots & 0 \\ 0 & 0 & 0 & \frac{1}{\mu_n} \end{bmatrix}$$

$$\mathbf{D}^{-1}\mathbf{V}_{\mu}\mathbf{D}^{-1} = \begin{bmatrix} \frac{1}{\phi} + \frac{1}{\mu_1} & 0 & 0 & 0 \\ 0 & \frac{1}{\phi} + \frac{1}{\mu_2} & 0 & 0 \\ 0 & 0 & \ddots & 0 \\ 0 & 0 & 0 & \frac{1}{\phi} + \frac{1}{\mu_n} \end{bmatrix}$$

Which leads to

$$\mathbf{W^{-1}} = \begin{bmatrix} \phi + \mu_1 & 0 & 0 & 0 \\ 0 & \phi + \mu_2 & 0 & 0 \\ 0 & 0 & \ddots & 0 \\ 0 & 0 & 0 & \phi + \mu_n \end{bmatrix}.$$

Milo NB-GLMM benchmarking

We first benchmarked our NB-GLMM implementation against several commonly-used R GLMM packages (Methods). These include penalised quasi-likelihood (glmm-PQL) [6], penalised iteratively reweighted least squares (glmer) [7] and auto-differentiation (glmmTMB) [8]. The latter is notable for its speed and accuracy and can be considered a gold standard. We simulated counts from a negative binomial distribution, allowing for additional variation based on fixed and random effect variables (Methods). Milo2.0 was able to accurately estimate the fixed effect and variance parameters across a range of ground truth values (Supplementary Figure 2). We further compared the performance of our NB-GLMM using simulated data from genetically related individuals (Methods) using either family membership or the full kinship matrix for the random effect. Similar results were obtained using glmmTMB, PQL and Milo2.0 with the family ID as the random effect variable. Using the kinship matrix as input to our NB-GLMM yielded similar results (Supplementary Figure 3), demonstrating the utility of this additional capability of Milo2.0. In sum, these benchmark analyses demonstrate that our NB-GLMM implementation is accurate and robust when applied to simulated data.
