## Supplemental Figures for "Milo2.0 unlocks population genetic analyses of cell state abundance using a count-based mixed model"

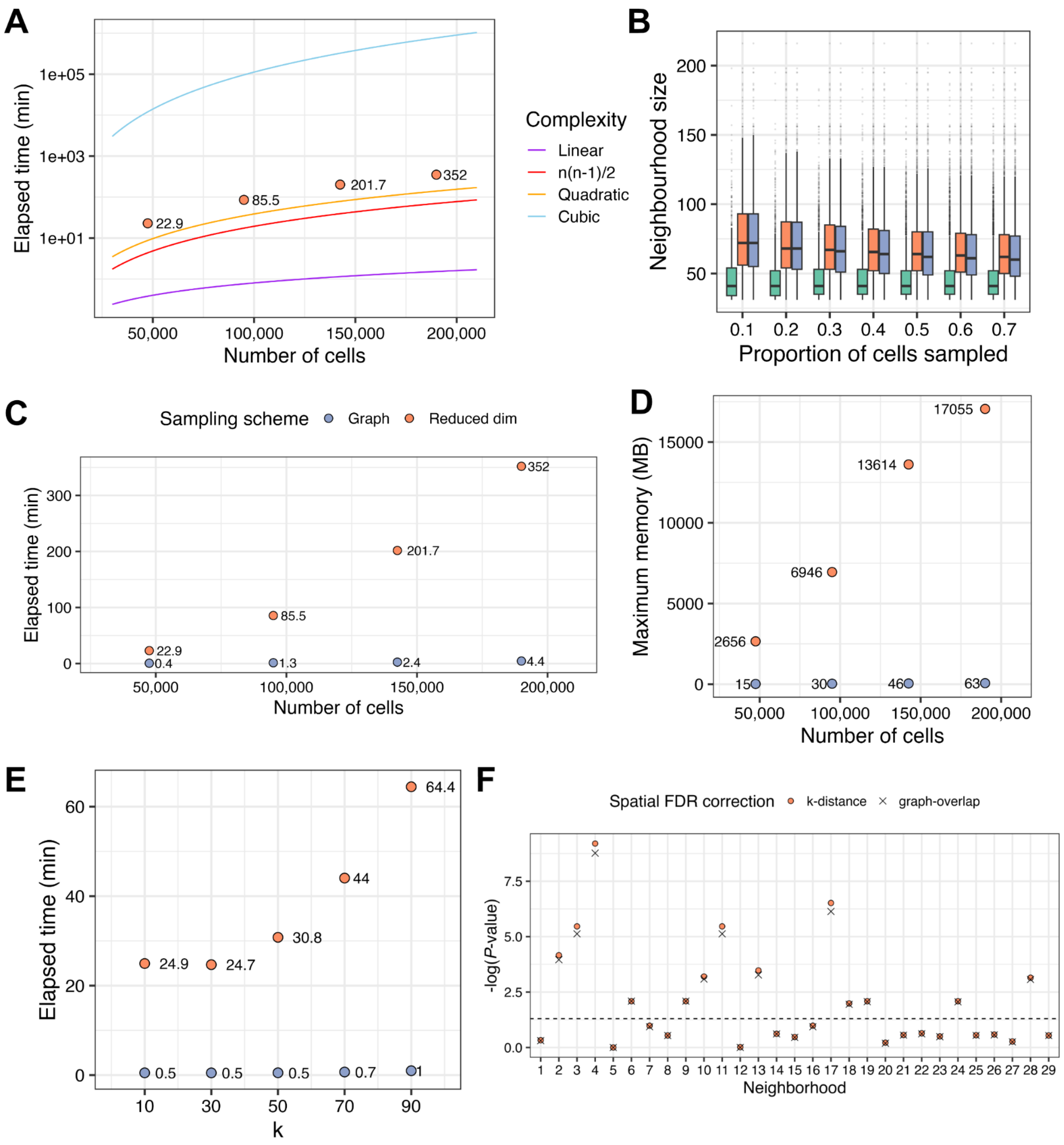


**Supplementary Figure 1.** (A) Computational load of Milo workflow. Coloured lines denote theoretical scaling with larger numbers of single-cells. Orange denotes show the elapsed time required to complete the Milo workflow from neighbourhood refinement to differential abundance testing. Y-axis shows the compute time (log scale), and x-axis the number of simulated single cells. (B) Neighbourhood sizes across a simulated data set after neighbourhood refinement using distance-based (orange) or graph-based (purple) algorithms. Uniform sampling is shown in the green boxes. Boxplots denote the median and interquartile range. Whiskers extend 1.5x IQR with outliers shown as points beyond this range. (C) Elapsed time (minutes) comparing distance(orange) and graph-based (purple) neighbourhood refinement algorithms. (D) Memory usage comparing distance (orange) and graph-based (purple) neighbourhood refinement algorithms. (E) The impact of varying k parameter on elapsed time for distance (orange) and graph-based (purple) neighbourhood refinement algorithms. (F) FDR correction is unbiased by graph-based approximation to the spatial FDR correction. Shown are $-\log_{10}$ spatial FDR values computed with distances (orange dots) or neighbourhood overlap (crosses) for 29 simulated neighbourhoods.

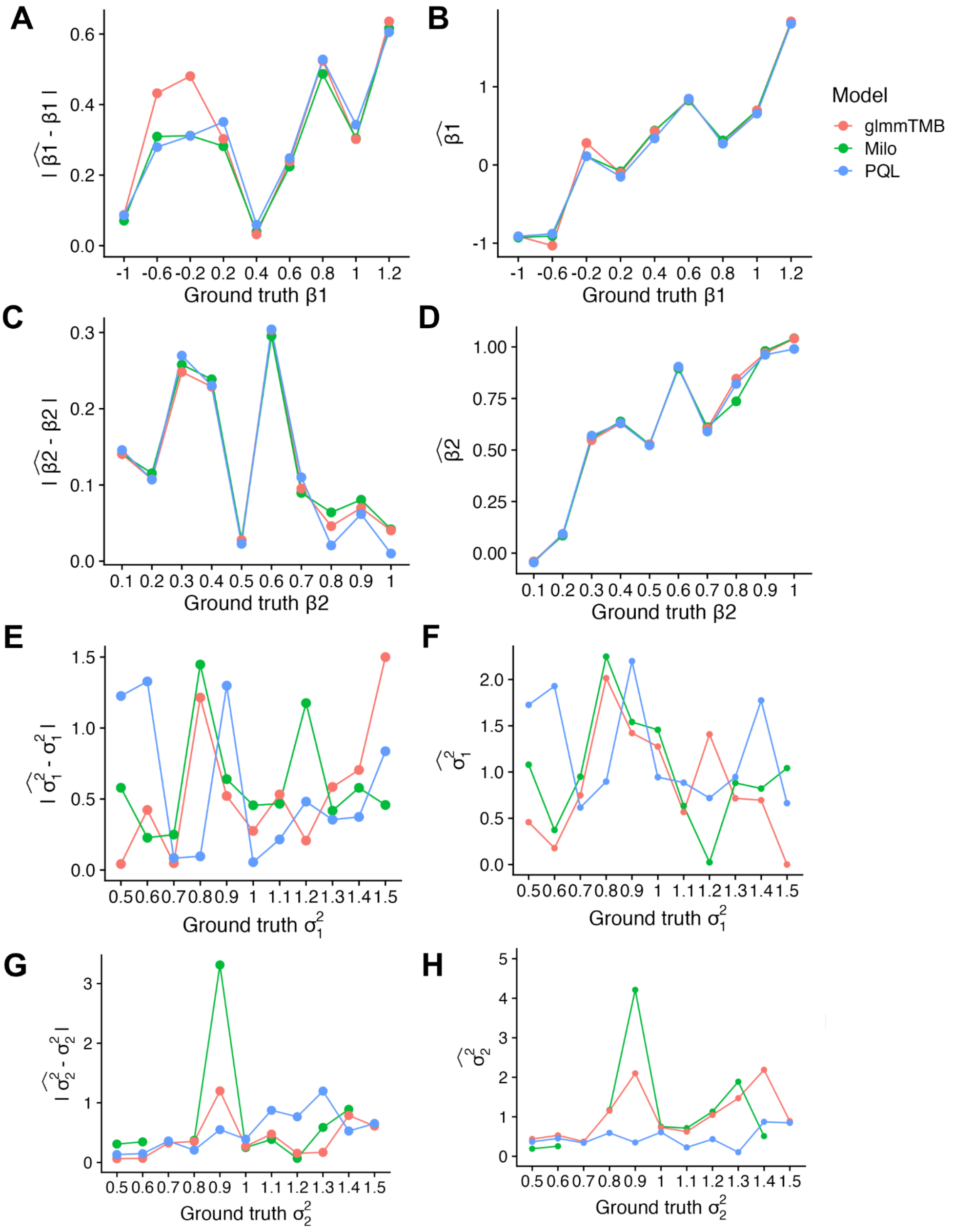


**Supplementary Figure 2.** Absolute deviation of parameter estimates for simulated parameters for fixed effect (A, C) and random effect variables (E, G). Comparisons of estimates to parameter ground truth values for fixed effect (B, D) and random effect variables (F, H). Coloured lines denote estimates from Milo (green), glmmTMB (red) or PQL (blue).

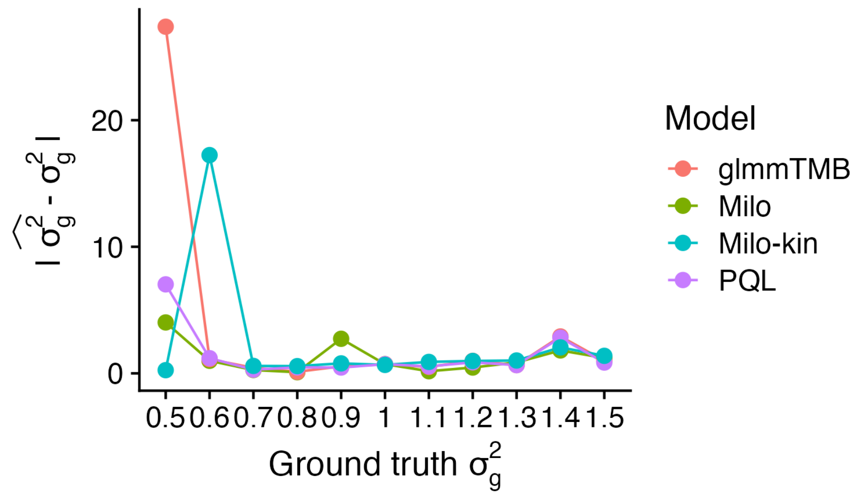


**Supplementary Figure 3.** Variance parameter estimate deviations from simulated related individuals. Input uses either family ID as an indicator random effect variable (glmmTMB, Milo, PQL) or a kinship covariance matrix (Milo-kin).

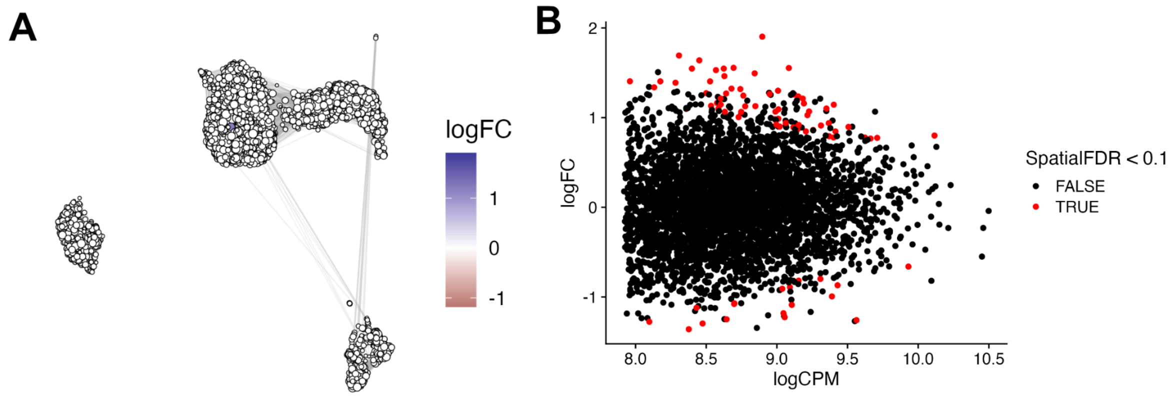


**Supplementary Figure 4.** Milo GLM DA testing results comparing AFR vs. EUR-ancestry similar individuals adjusting for batch as a fixed effect variable. (A) Log fold change overlaid on the UMAP as in Figure 1. (B) MA plot denoting log fold changes across neighbourhood abundance. Points are coloured red if they are statistically significantly DA (FDR 10%).

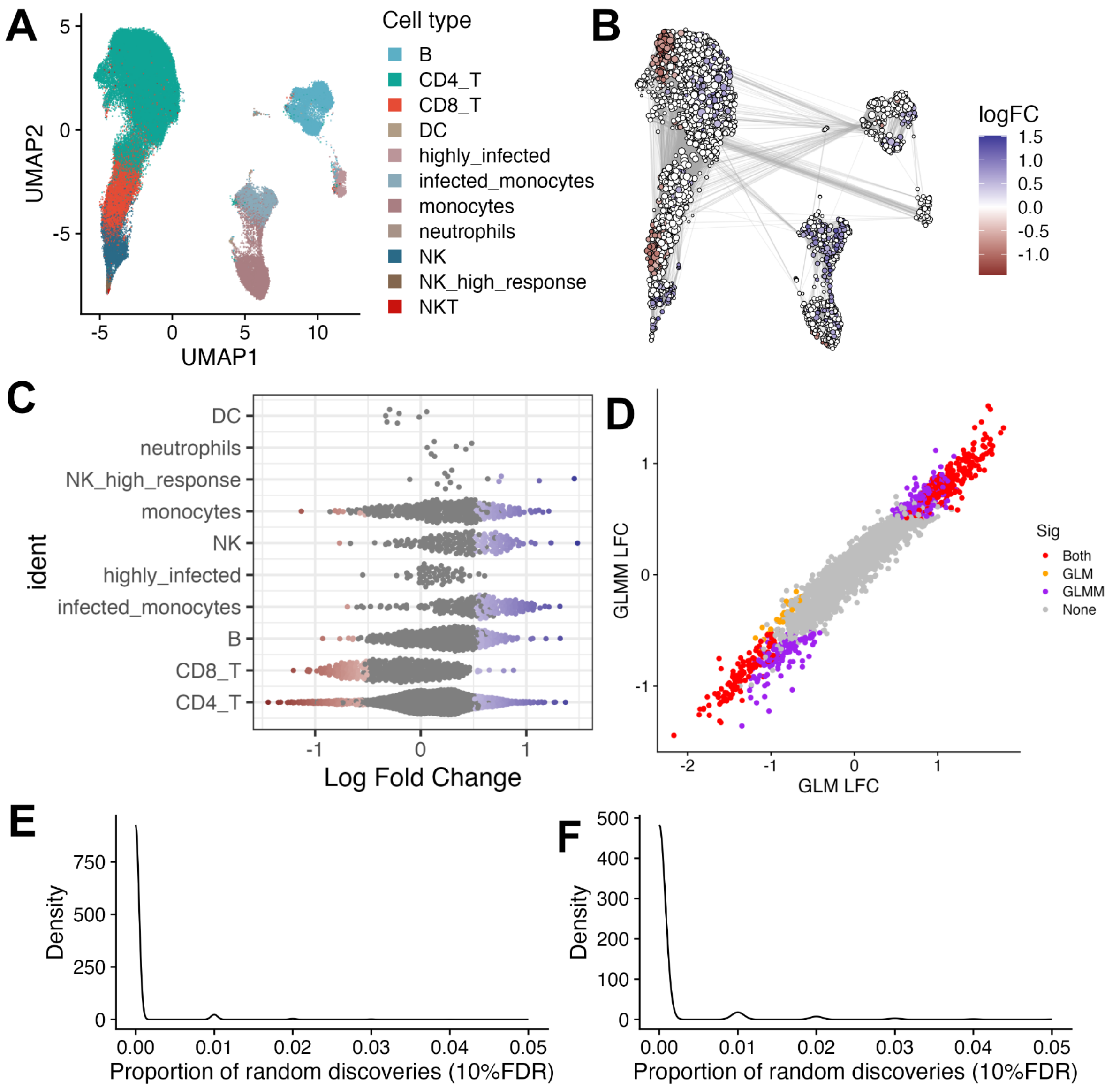


**Supplementary Figure 5.** (A) UMAP of IAV-infected PBMCs from Randolph *et al.* Single-cells are coloured by the cell type label from the original study. (B) DA results as log fold change (logFC) overlaid on the neighbourhood UMAP, where each point denotes a neighbourhood of cells anchored to the UMAP co-ordinates in (A) by the index cell. DA results are from comparing AFR vs. EUR-ancestry similar individuals. (C) DA results of AFR vs. EUR-ancestry similar individual results (x-axis, log fold change) compared to the cell type label annotations as in (A), highlighting the heterogeneity in effect sizes and directions across peripheral immune cell types. (D) Comparison of GLMM (y-axis) vs. GLM (x-axis) log fold change estimates illustrates the concordance between models. Points are neighbourhoods coloured by whether they are statistically DA (FDR 10%) in the GLM (orange), GLMM (purple), both (red) or neither (grey). (E-F) Density plots of randomly shuffled mock (E) and IAV infected (F) spatial FDR values.

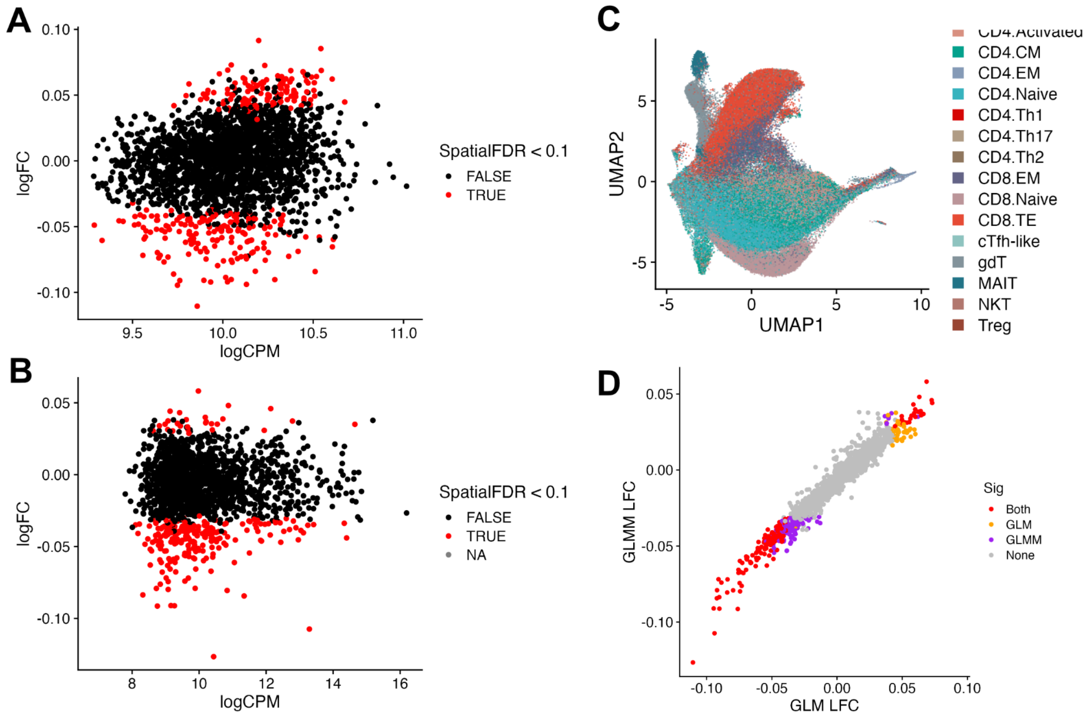


**Supplementary Figure 6.** MA plots of DA results of COVID19 severity, adjusting for batch as a fixed effect (A) or random effect variable (B). (C) UMAP denoting batch-corrected T cell data, with points coloured by original cell type labels from Stephenson *et al.* (D) Concordance of DA results from the GLMM (y-axis) and GLM (x-axis). Neighbourhood points are coloured by whether they are statistically significant (FDR 10%) in the GLM (orange), GLMM (purple), both (red) or neigher (grey).

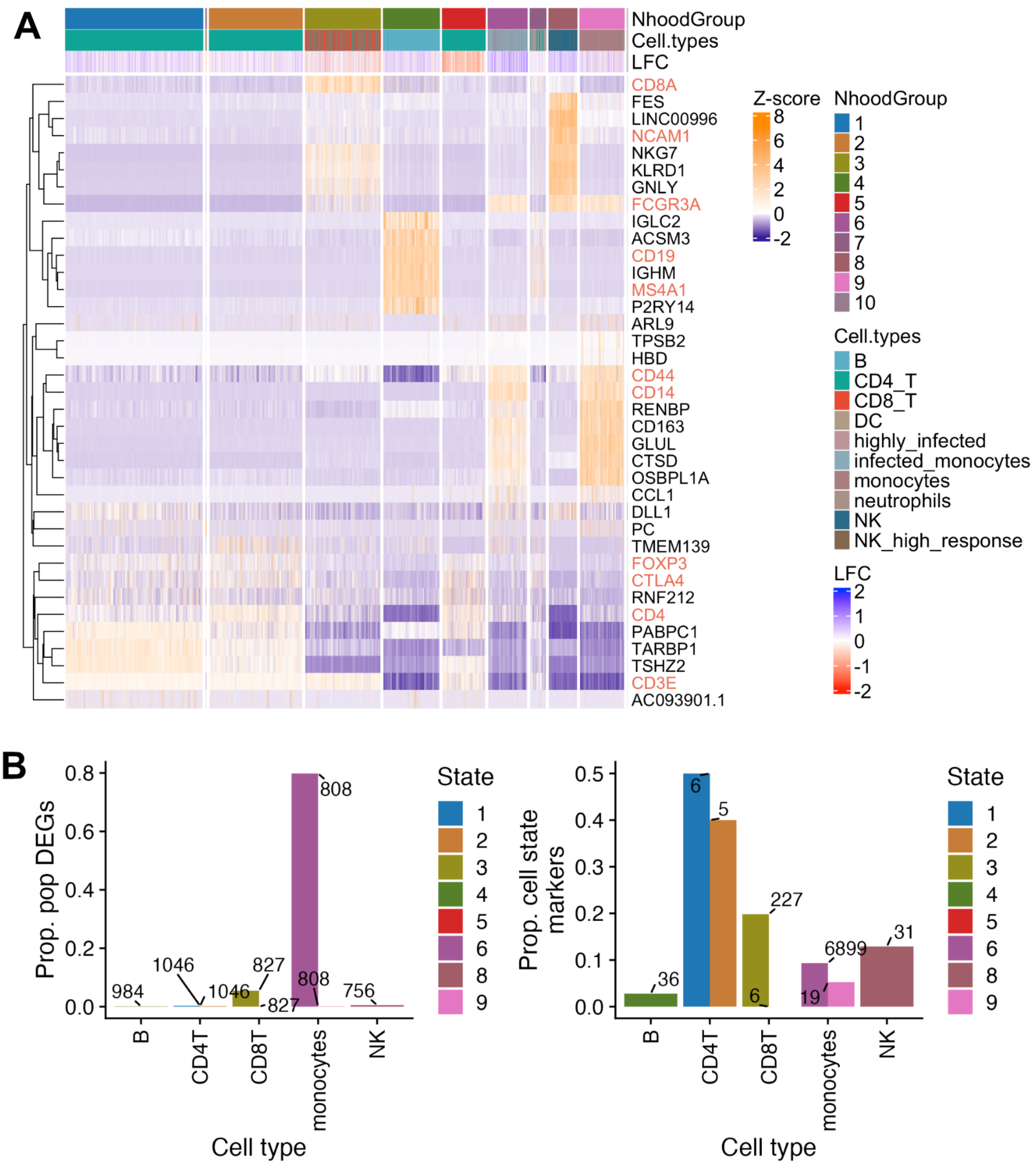


**Supplementary Figure 7.** (A) Heatmap of IAV-infected neighbourhood group top marker genes. Gene expression is shown as Z-scores across neighbourhoods (columns). Highlighted genes (rows) denote canonical immune cell type marker genes. (B) Proportions of popDE genes (left) and popDA cell state markers (right) explained by the respective gene set form IAV infected peripheral immune cells.

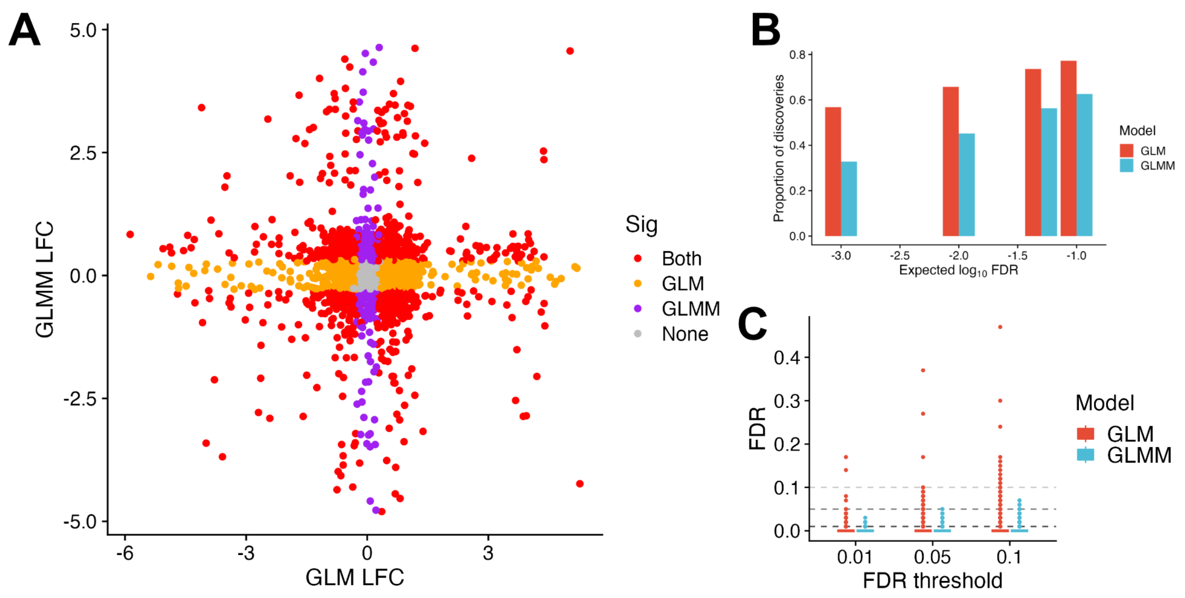


**Supplementary Figure 8.** (A) Comparison of log fold change from GLMM (y-axis) and GLM (x-axis) using repeated measures for each individual. Points denote neighbourhoods identified as DA (FDR 10%) in the GLM (orange), GLMM (purple), both (red) or neither analysis (grey). (B) The proportion of discoveries at different FDR thresholds. X-axis shows the -$\log_{10}$ spatial FDR. Bars are coloured by the GLM (red) and GLMM (blue). (C) The proportion of false discoveries (FDR, y-axis) across 100 randomly shuffled data sets analysed with the GLM (red) or GLMM (blue).

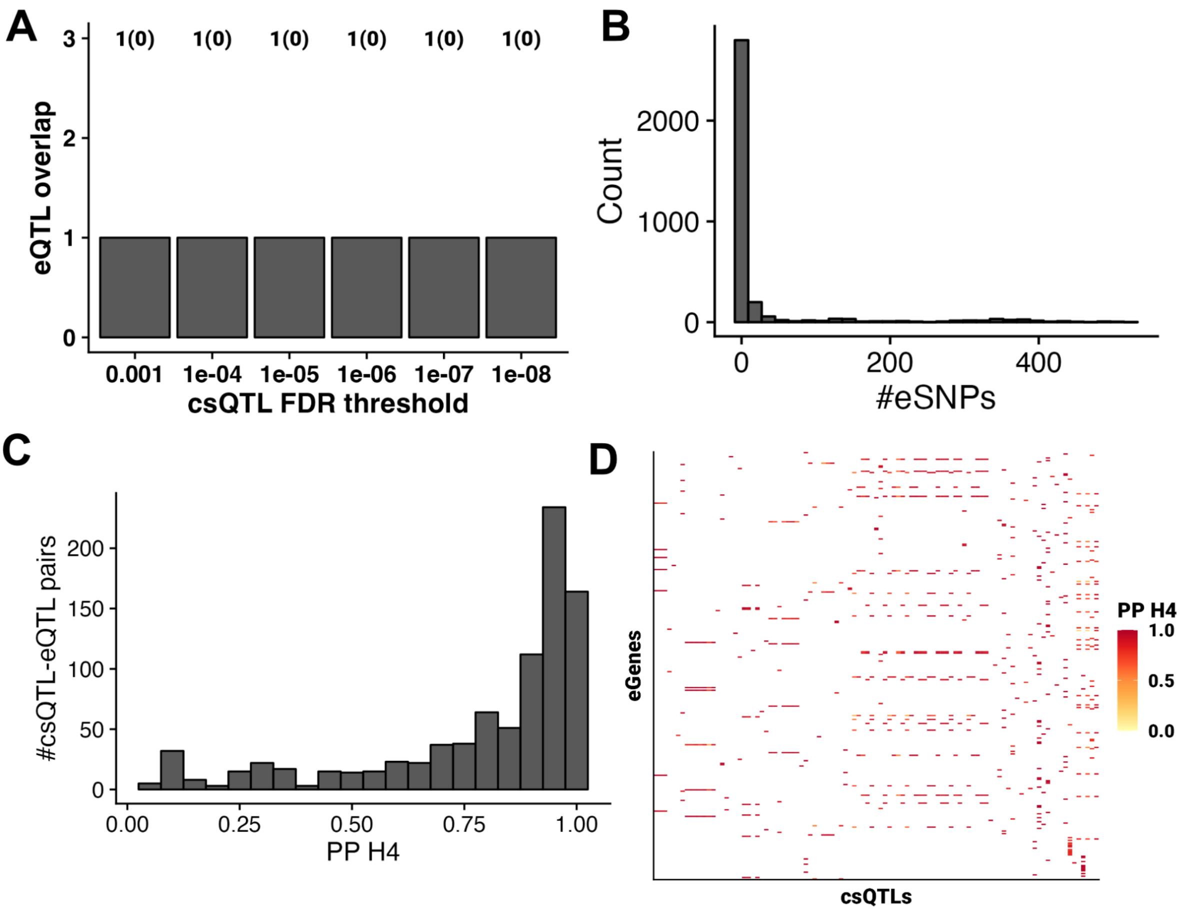


**Supplementary Figure 9.** (A) Randolph *et al.* eQTL overlap with csQTLs at increasing FDR thresholds. (B) Histogram of the number of eSNPs at each csQTL for the eQTLs tested. (C) Histogram of H4 posterior probabilities highlights the enrichment of colocalised neighbourhood eQTLs with csQTLs. (D) Heatmap of eGenes linked to csQTLs coloured by the H4 posterior probability. csQTLs (columns) are sorted according to chromosome and genome position.

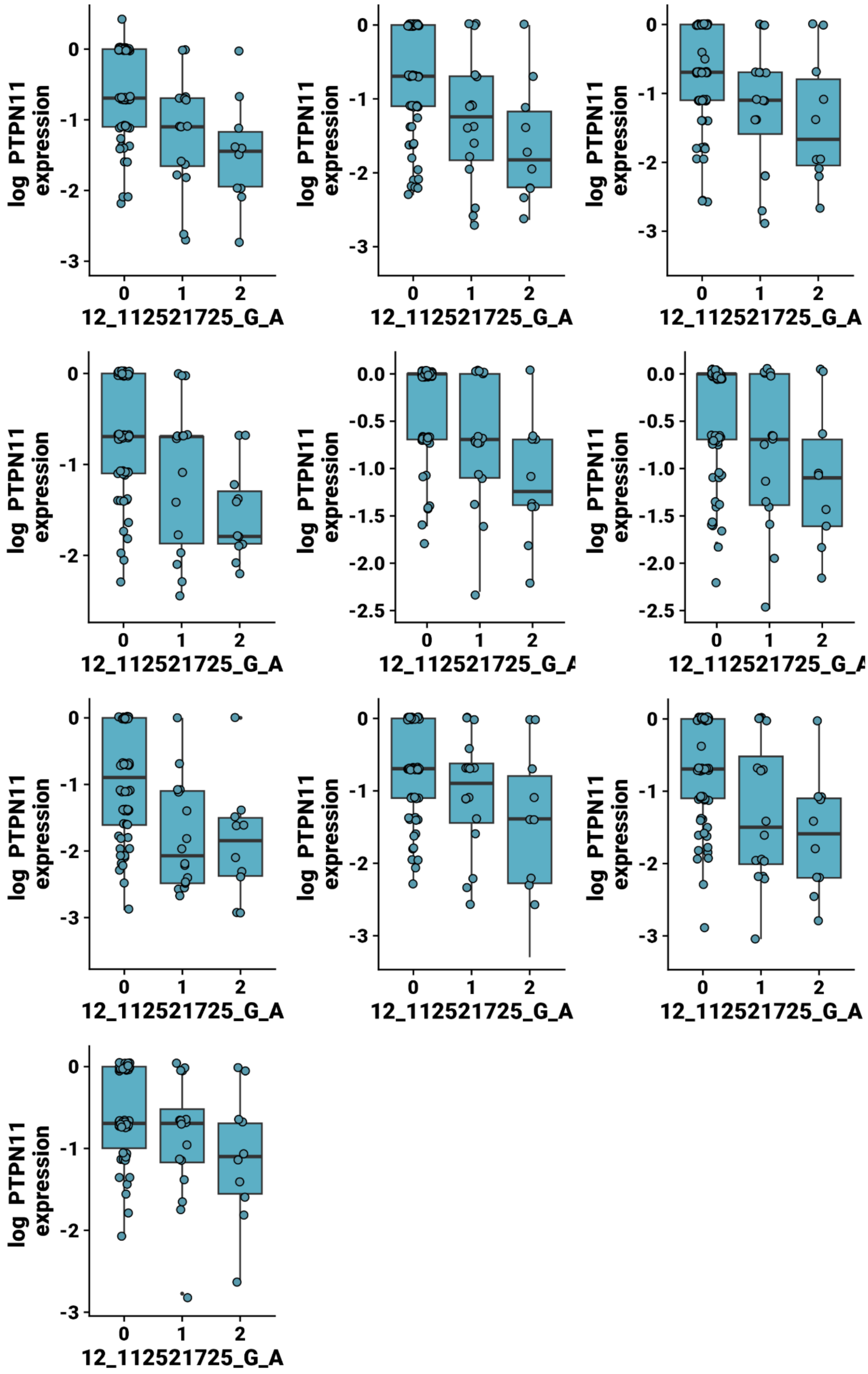


**Supplementary Figure 10.** Box and scatter plots showing *PTPN11* expression across n=89 samples by genotype at the SNP 12_112521725_G_A. Points represent the average expression across cells for each individual for each of the neighbourhoods shown. Boxes are the median and interquartile range, with whiskers extending 1.5x the IQR.

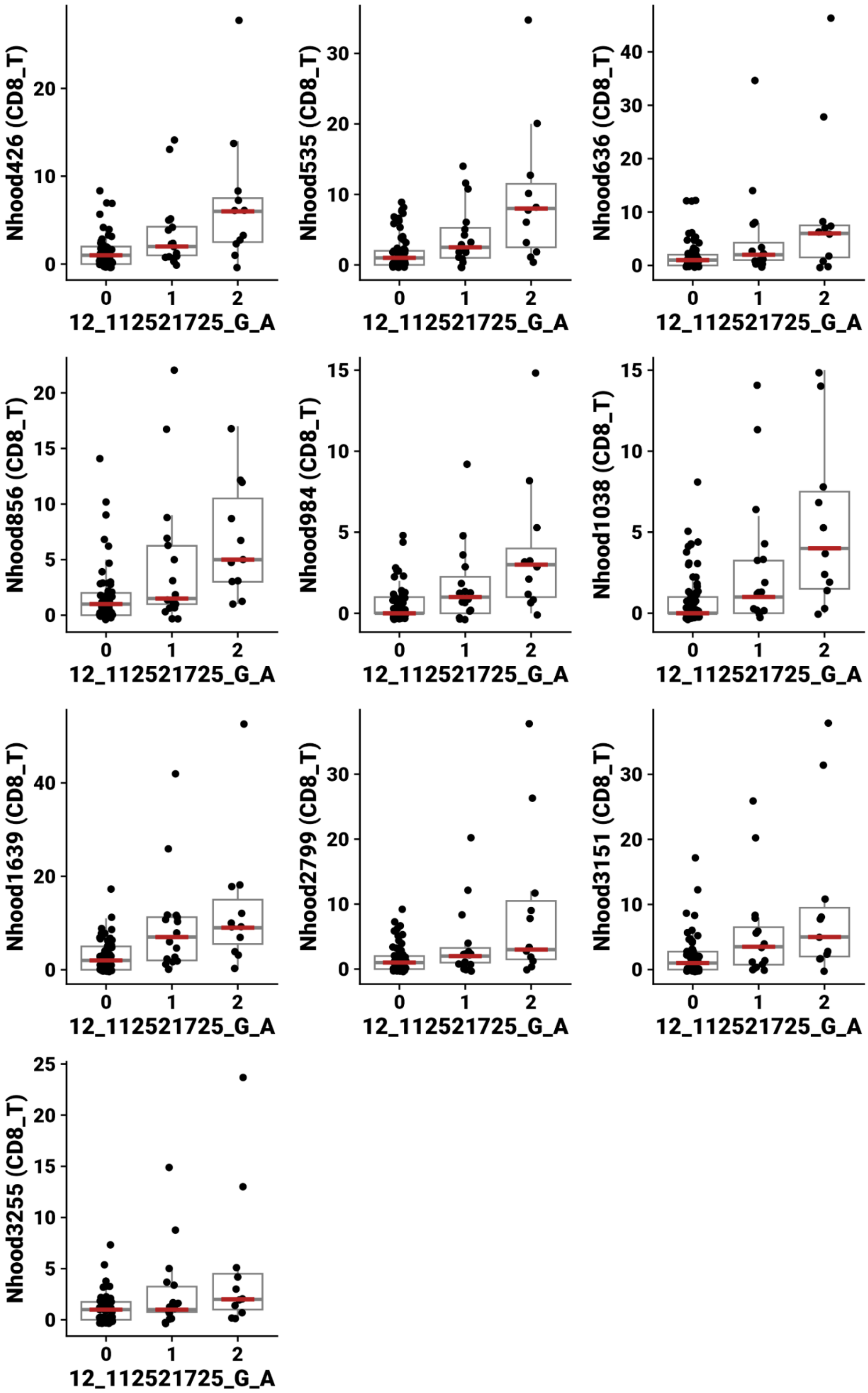


**Supplementary Figure 11.** Box and scatter plots showing neighbourhood abundance across n=89 samples by genotype at the SNP 12_112521725_G_A. Points represent the number of cells for each individual for each of the neighbourhoods shown. Boxes are the median and interquartile range, with whiskers extending 1.5x the IQR. The order of boxplots is the same as for Supplementary Figure 10.

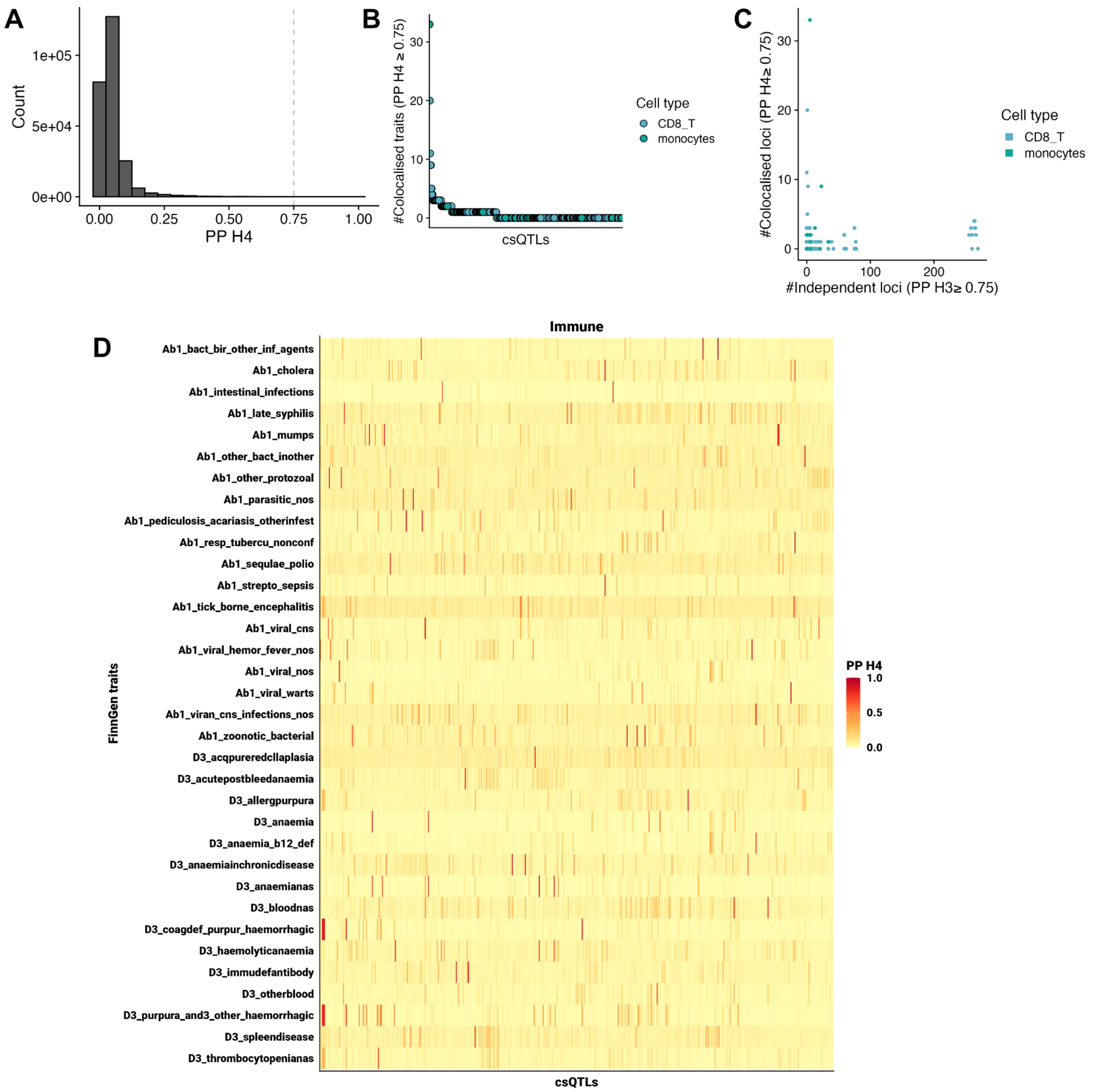


**Supplementary Figure 12.** (A) Distribution of colocalisation H4 posterior probabilities between csQTLs and FinnGen traits. (B) Ranked csQTLs but the number of traits with strong evidence (H4 PP ≥ 0.75) of colocalisation with FinnGen study traits. (C) The relationship between colocalisation (y-axis; H4 PP) and independent signals (x-axis; H3 PP) for csQTLs and FinnGen study traits. Points in (B) and (C) are coloured by the cell type annotation to which the neighbourhood corresponds, as in Figure 2A. (D) Heatmap of immune system related traits from the FinnGen study that show colocalisation with at least one csQTL.

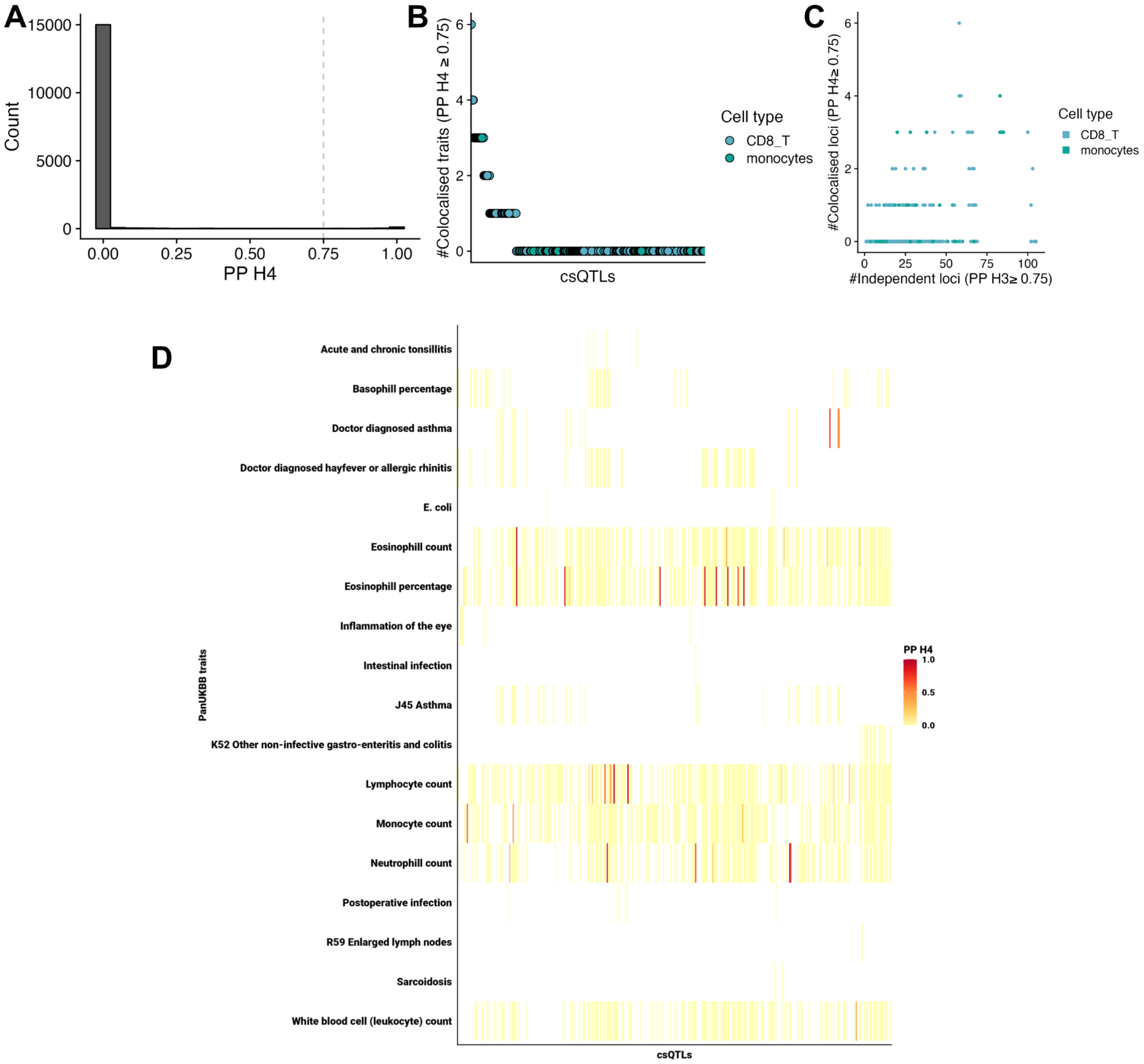


**Supplementary Figure 13.** (A) Distribution of colocalisation H4 posterior probabilities between csQTLs and PanUKBB traits. (B) Ranked csQTLs but the number of traits with strong evidence (H4 PP ≥ 0.75) of colocalisation with PanUKBB study traits. (C) The relationship between colocalisation (y-axis; H4 PP) and independent signals (x-axis; H3 PP) for csQTLs and PanUKBB study traits. Points in (B) and (C) are coloured by the cell type annotation to which the neighbourhood corresponds, as in Figure 2A. (D) Heatmap of immune system related traits from the PanUKBB study that show colocalisation with at least one csQTL.

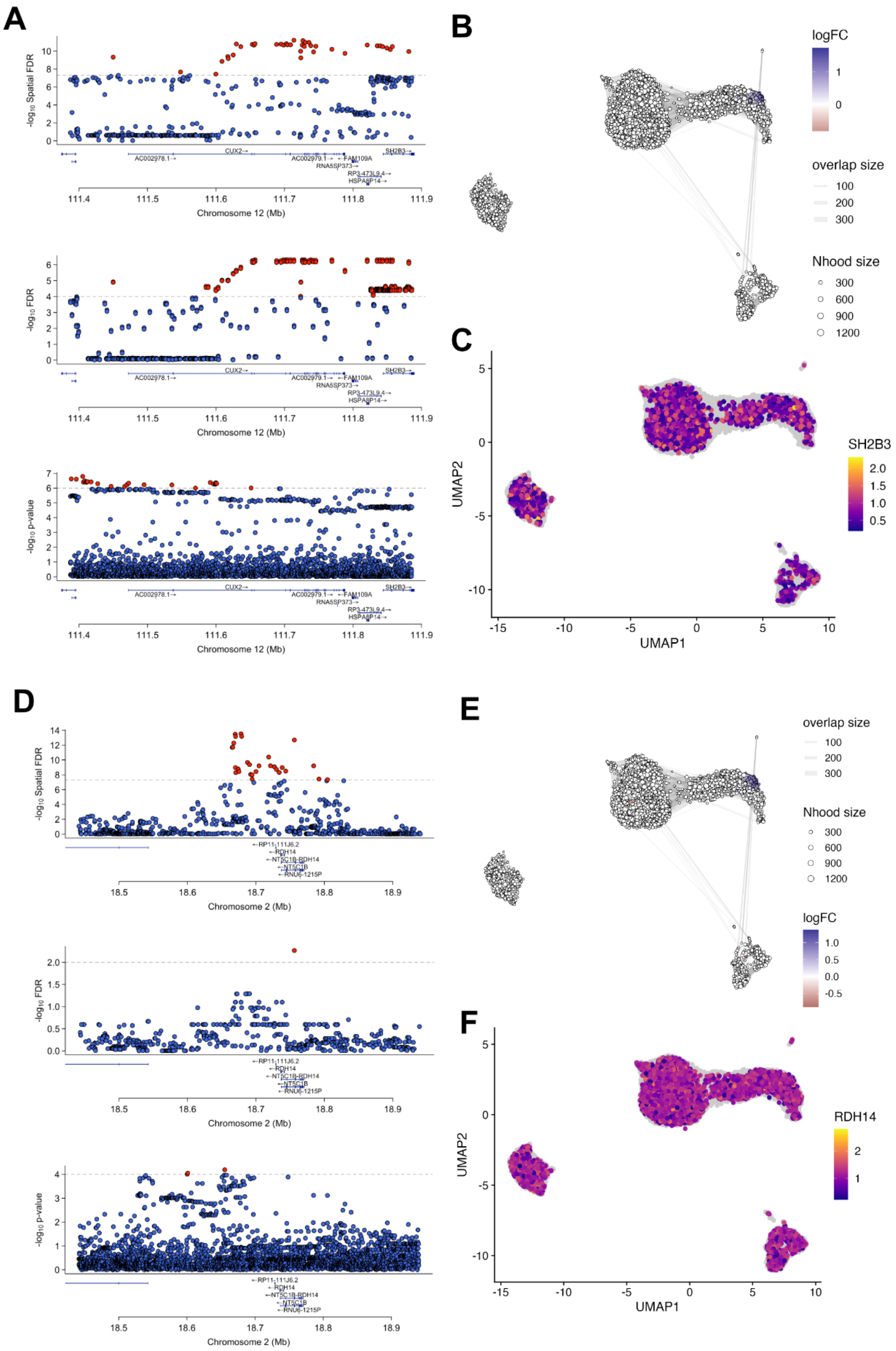


**Supplementary Figure 14.** (A) LocusZoom plots showing the shared genetic signal on chromosome 2 for a CD8 T cell csQTL (top), *SH2B3* eQTL (middle) and systemic connective tissue diseases from the FinnGen studyy (bottom). (B) csQTL log fold change overlaid on the neighbourhood UMAP as in Figure 2A-B. (C) *SH2B3* expression across single cells overlaid on the UMAP as in Figure 2A. (D) LocusZoom plots of the shared genetic signal on chromosome 2 for a CD8 T cell csQTL (top), *RDH14* eQTL (middle) and mumps from the FinnGen study (bottom). (E) csQTL log fold change for the csQTL in (D-top) overlaid on the neighbourhood UMAP, as in Figure 2A-B. (F) *RDH14* expression across single cells overlaid on the UMAP as in Figure 2A.

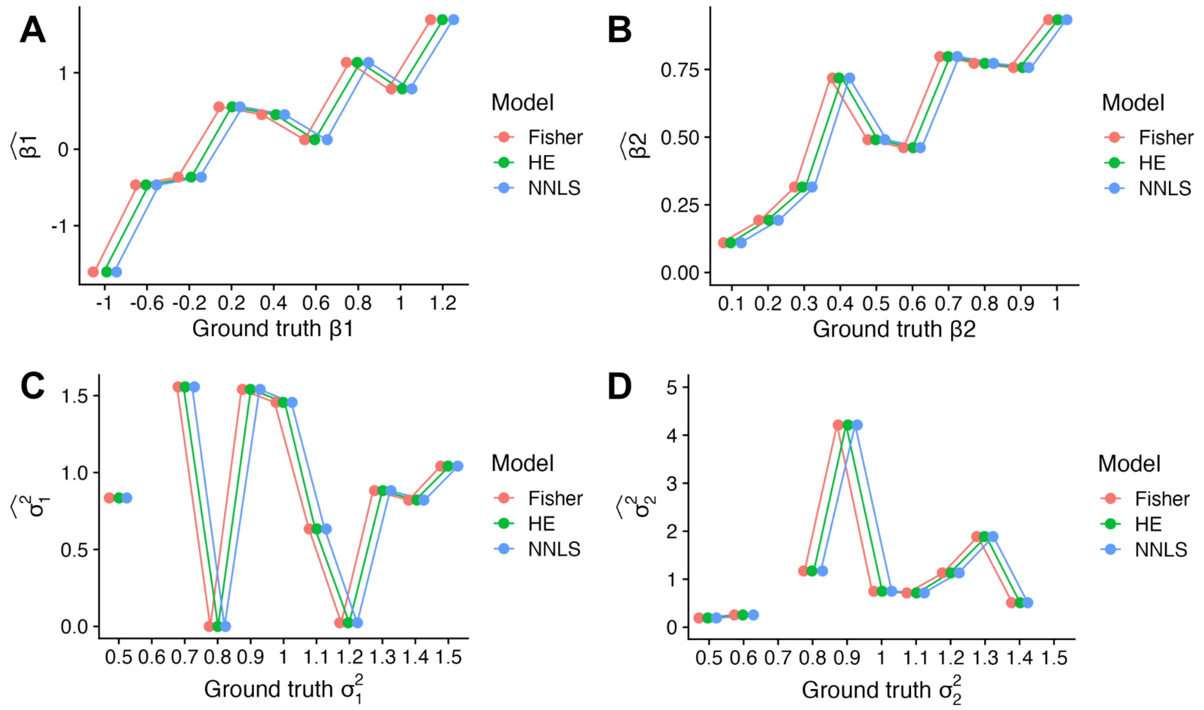


**Supplementary Figure 15.** (A-B) Fixed effect parameter estimates on simulated unrelated individuals for a single neighbourhood (n=150). Points denote the estimates (y-axis) compared to the ground truth values. (C-D) Variance parameter estimates on the same simulated data. Missing data points reflect failure modes of the model. Points are coloured by the estimator: Fisher (salmon) or Haseman-Elston (turquoise). Points are jittered on the x-axis to illustrate the estimates are equivalent.
